# Cognitive dysfunction and anxiety resulting from synaptic downscaling, hippocampal atrophy and ventricular enlargement with intracerebroventricular streptozotocin injection in male Wistar rats

**DOI:** 10.1101/2022.04.04.486747

**Authors:** Avishek Roy, Sakshi Sharma, Tapas Chandra Nag, Jatinder Katyal, Yogendra Kumar Gupta, Suman Jain

## Abstract

Insulin-resistant brain state is proposed to be the early sign of Alzheimer’s disease(AD), which can be studied in intracerebroventricular streptozotocin (ICV-STZ) rodent model. ICV-STZ is reported to induce sporadic AD with the majority of the disease hallmarks as phenotype. On the other hand, Available experimental evidence has used varying doses of STZ (<1 to 3mg/kg) and studied its effect for different study durations, ranging from 14-270 days. Though these studies suggest 3mg/kg of ICV-STZ to be the optimum dose for progressive pathogenesis, the reason for such is elusive.

Here we sought to investigate the mechanism of action of 3mg/kg ICV-STZ on cognitive and non-cognitive aspects at a follow-up interval of two weeks for two months. On 60th day, we examined layer thickness, cell density, ventricular volume, spine density, protein expression related to brain metabolism and mitochondrial function by histological examination. Findings suggest progressive loss of spatial, episodic, avoidance memory with increase in anxiety in a span of two month. Furthermore, hippocampal neurodegeneration, ventricular enlargement, diffused amyloid plaque deposition, loss of spine in dentate gyrus and imbalance in energy homeostasis was found on 60th day post injection. Interestingly, AD rats showed a uniform fraction of time spent in four quadrants of water maze with change in strategy when they were exposed to height. Our findings reveal that ICV-STZ injection at a dose of 3mg/kg can cause cognitive and neuropsychiatric abnormalities due to structural loss both at neuronal as well as synaptic level, which is tightly associated with change in neuronal metabolism.

## Introduction

AD is one of the commonest forms of dementia, and characterized by the deposition of hyperphosphorylated tau, amyloid beta (Aß) in cerebral parenchyma. These proteopathies are tightly associated with cognitive impairment. Cognitive impairment in AD is reported to be resulted from synaptic dysfunction and loss of adult neurogenesis. On the other hand, insulin resistance brain state (IRBS) is the gold standard for early detection of AD in patients using FDG-PET imaging. Streptozotocin (STZ) is a glucosamine nitrosourea compound, which if injected into the brain intracerebroventricularly (ICV) produces IRBS condition resembling sporadic AD over a period of time. Effect of varied doses (1.5-3mg/kg, single/multiple injections) of STZ was studied for short (14-21days), mild (30-42 days), moderate (90-105 days) and long (250-170 days) time period for pathological events. Those studies indicated 3mg/kg to be the best choice for induction of the sporadic AD in rodents, in terms of cognitive decline. And this was associated with many micro and macro structural changes ranging from mitochondrial dysfunction to cortical thinning. However, relevant studies were not available deciphering the morpho-functional pathological processes after 3mg/kg ICV-STZ injection. Previous studies with different time-periods have reported that short duration treatment leads to cognitive impairment, anxiety, hp-tau deposition, neurodegeneration, and gliosis on 14th day, whereas GSK3ß increment occurs in the hippocampus on 30th day after STZ injection (Salkovic-Petrisic et al. 2006; Chen et al. 2013). These phenotypes were observed due to cellular, mitochondrial and insulin dysfunction of brain (C. Correia et al. 2013). However, observation for moderate time period revealed increase in Aß1-42 and amyloid plaque deposition, expression of AKT, insulin1 and 2 mRNA and insulin receptor in the hippocampus and frontoparietal cortex on the 80-90th day with a trend for reversal from acute cognitive impairment on 14th day (Salkovic-Petrisic et al. 2006; Grünblatt et al. 2007; Rostami et al. 2017). Long temporal observation have shown disturbance of avoidance memory following a biphasic pattern, with insult to the parietal cortex and corpus callosum (Knezovic et al. 2015). On the basis of these studies some authors tested novel drugs against AD, viz. antia (El Sayed and Ghoneum 2020), PPAR agonist (Lin et al. 2022), QTC-4-MeOBnE (Fronza et al. 2019), magnesium (Xu et al. 2014). When ICV-STZ has been compared to 3xTg model of AD, results showed altered expression of 84 genes related to APP processing, apoptosis, mTOR and insulin signaling and impairments in episodic memory, anxiety, but no change in motor activity or related postural balance after unilateral injection of 3mg/kg ICV-STZ in female mice at 42 day post injection (Chen et al. 2012, 2013). In a recent study, Bao et al. (2017) reported gender-specific differences in the effect of ICV-STZ in cognitive as well as neuronal morphometry in Sprague-Dawley rats, wherein females responded less to the STZ toxicity in comparison to males, due to protective role in estradiol in the former (Bao et al. 2017). More recently, Gáspár et al. (2021) reported that long Evans rats are less sensitive to ICV-STZ insults than Wistar, suggesting a strain diversity in this model (Gáspár et al. 2021).

Therefore, in the present study, we had used male Wistar rats to decipher the pathological pattern of disease by means of behavioral and morphological changes at 3mg/kg (ICV, single injection, STZ). We have first followed up the spatial, episodic, avoidance memory and anxiety level simultaneously at an interval of 15 days until two months. At the end of the study period, we have examined hippocampus and its associated structures for cell density, volume analysis on cresyl violet stained coronal sections and morphometry on Golgi-Cox stained coronal sections of brain. We measured expression of markers viz. GSK3ß, PI3K, mt-COX1 in limbic system and also looked for amyloid plaques using Congored staining. Finally, apart from the regular analysis of behavioral data, we further looked at the distribution of fraction of time spent on 60th day probe trial in Morris water maze (MWM) and ethograms of animals on 60th day in elevated plus maze (EPM) in order to dissect the functional changes in cognitive and neuropsychiatric domain. Our findings indicates that ICV-STZ injection causes progressive loss of retention of memory in all the domains with gradual increase in anxiety. Interestingly, our data showed that on 60^th^ day post injection of STZ rats spent uniform time in all four quadrants of water maze. Whereas, they change the exploration strategy evident by ethogram analysis in elevated plus maze more specifically at the initial moment of decision animals were in stretch-attend posture. These cognitive and neuropsychiatric load may be due to the neurodegeneration, ventricular atrophy, amyloid plaque deposition and loss of spine density. Immunofluorescence examination at limbic system revealed that these multimodal changes are due to abnormality in expression of proteins related to insulin signaling and mitochondrial physiology. Hence this study confirms the cause of morpho-functional effects of ICV-STZ (3mg/kg) reported in previous studies.

## Materials and Methods

### Animals

Total 54 male Wistar rats (280 ±10 gm) were included in the present study, which were maintained in standard housing conditions at an ambient room temperature of 25±1°C, relative humidity of 50-55% and the light-dark cycle of 14:10 h. Rats were provided with *ad libitum* access to food and water, each housed individually in polypropylene cages (50 cm X 20 cm X15 cm) with metallic lid, which has separate laboratory chow and water bottle holders. All precautions were taken during the period of housing to avoid pain. All experimental protocols were approved by institutional animal ethical committee (*file no. 937/IAEC/PhD-2016*), in accordance with committee for the purpose of control and supervision of experiments on animals (CPSCEA) guidelines.

Rats were randomly divided into three groups viz. Control, Sham and AD, where Sham and AD group of animals were injected with artificial cerebrospinal fluid (aCSF) or STZ, respectively in the lateral ventricles. Among the 18 rats recruited in each group, 16, 15 and 12 rats were survived till the end of the study period in Control, Sham and AD group, respectively. Therefore, total survival rate in all experimental groups is 68%; furthermore, mortality rate in 3mg/kg STZ injected rats were up to 33.33%. For behavioral experiments, we used ≥9 animals, for histological and biochemical experiments we recruited 3 and 4 animals/group.

### Experimental design

After adaptation to the departmental animal facility, rats were randomly divided into three groups using block-size randomization (Efird 2011):

Control: age matched cage control animals housed in same condition to others

Sham: 5µl/site bolus injection of aCSF bilaterally at lateral ventricles

AD: 3mg/kg of b/w bolus injection of STZ bilaterally at lateral ventricles

Primarily, all baseline behavior was recorded for all experimental groups within first week after adaptation period to the colony room. Next, groups were homogenized based on baseline behavioral data in order to avoid any baseline bias. Animals were undergone surgery for Sham/ AD group with aCSF/STZ, respectively. From then animals were tested for their cognitive and non-cognitive behaviors at an interval of 15 days for a stretch of two months. At the end of the study period, animals were sacrificed either via transcardial perfusion or CO_2_ concussion for histological and biochemical experiments, respectively.

### Behavioral assessments

In present study, MWM, one trial object recognition (OTORT), step-down passive avoidance (SDPA) and EPM tests were chosen for spatial reference, episodic recognition, avoidance memory and unconditioned anxiety due to fear of height, respectively. All behavioral assessments were recorded and analyzed with video tracking devices except for passive avoidance test, which was conducted manually. Behavioral execution in cognitive and non-cognitive tasks were recorded on 15^th^, 30^th^, 45^th^ and 60^th^ day post-injection of aCSF/STZ accordingly. The order of tests was OTORT, EPM, SDPA, MWM with interval 50±10 minute to avoid any effect within them, and it remained same for every follow-up days (fig.1. panel 1).

### MWM test

It was conducted to decipher reference spatial memory in the experimental animals as described previously (Morris 1984; Mehla et al. 2012a). In MWM a water tank (metallic, customized, 170cm dia. X 72cm height) has been divided into four equal imaginary quadrants with the help of video-tracking software. Animals were trained to remember how to escape swimming by entering onto hidden platform (10 cm dia. X 35 cm height). The hidden platform was situated in fourth quadrant for two minutes in consecutive four days, where fourth quadrant was refereed with an intra-maze cue. A training trial was started by placing the animal facing towards wall of the tank. Trial was terminated once animal sit on the platform for 10 or 120 sec elapsed, whichever condition meet first. If animals did not reach destiny on time, they were manually guided to the platform by the experimenter.

On the fifth day, a spatial probe trial was conducted where protocol was same as mentioned in previous paragraph except animals were allowed to explore the tank only for 60 sec and no platform has been placed inside the tank. All trials have been recorded and data acquired using AnyMaze software (ver.05; Stoelting Corp., USA). Furthermore, only probe trial was repeated on the follow-up days post-injection for retention of spatial memory.

### Dirichlet distribution of swimming strategy in MWM

Dirichlet distribution is a multivariate generalization of beta distributions, where in the recent paper it has been used a stand-alone test to determine whether the animals spend the same time in the four imaginary quadrants of MWM tank (Maugard et al. 2019). Four quadrants were conceptualized as one target quadrant (TQ), one opposite quadrant (OQ), and two alternative quadrants (AQ). Null hypothesis was adapted only when a group’s fraction of time spent in TQ=AQ=OQ. Further, for understanding the outcome authors has proposed a new way to plot the data from the four quadrants as the aforementioned conceptions, where graph will show mean values and inter-individual variability at the same time, on an easily interpretable chart. For the analysis we used code available at https://github.com/xuod/dirichlet.

### One trial object recognition test

The open field was used to conduct one trial object recognition test according to (Ennaceur and Delacour 1988), which specifically target episodic memory. Wooden/metallic/glass/plastic objects with different shapes and sizes, they are of primary colors with different palpable textures were tested for their ‘interest index’ a parameter assessed for ability to induce interest for novelty (data in supplementary Fig.S1.a.); however, in the present study plastic/glass made objects were used.

In OTORT, animals were first habituated with an open field arena (50 cm X 50 cm, Coulbourne Instruments, USA) with an exploration time of 5 min two times/day for 3 days. On the fourth day, a ‘sample phase’ was conducted where animal was placed in the center of the arena facing towards wall with similar objects at a same distance from the wall (10 cm from the wall). With a delay of fifteen minutes of ‘sample phase’ animals were allowed to explore the same arena for 5min, except one of the sample object is being changed with a novel one. This phase is called ‘test phase’. We have chosen a time gap of 15 min between two phases to target short term memory. In both phases, exploration of the animals was tracked with CCD camera (Panasonic, Japan) using Ethovision XT (ver. 07, Noldus Co., Netherlands) and data acquired and analysed for recognition and discrimination index, using the formulae:

♦ Recognition index= {(time spent exploring novel object)-(time spent exploring familiar object)}/{(time spent exploring novel object)+(time spent exploring familiar object)}
♦ Discrimination index= {(time spent exploring novel object)-(time spent exploring familiar object)}/{(time spent exploring novel object)+(time spent exploring familiar object)}

On the follow-up time periods at an interval of 15days ‘test phase’ was repeated.

### Stepdown test for passive avoidance

Passive avoidance was tested in rats to assess long-term avoidance/fear memory deficits in accordance with (Filho et al. 2015). We have used modular animal test cage with metal grid floor which is part of Skinner Box (model: E10-10SF, Coulbourne Instruments, USA), this was connected with an animal test cage grid floor shocker (model: E13-08, Coulbourne Instruments, USA) with a safe bench/ wooden platform where animals were placed before starting the experiment. We have used ‘consolidation protocol’ consisting a training phase, and 24hr later a test phase. On training phase, rat was placed onto the safe bench and upon stepping down to grid floor with its four limbs it got an electric shock of 2 mA for 3 sec. Retention of memory was defined as avoiding foot shock by not being active on grid floor on second day/test phase. After 24 h of training phase same protocol was followed except foot shock upon step-down and behavior was observed for 3 min of time. In this observation, period number of step down (errors) and latency to the first step down on the grid floor was measured manually.

We repeated the second day/ test phase on every 15-day interval of post-injection of STZ in the experimental groups and same parameters were recorded as a measure of retention of fear memory.

### EPM test

EPM paradigm was conducted in order to decipher the anxiety like behavior, and this in accordance with previous method (Walf and Frye 2007). This behavior is based on the anxiety induced by the fear of height in rodents. We used an EPM setup consists of a customized plus maze with an opaque plastic board fixed on to iron frame made out of plywood. The length of the open and closed arm of 76 and 134 cm, respectively, with a junction of arms, i.e., central area of 6 × 6 cm. Animal’s behavior was video tracked with CCD camera (Panasonic, Japan) associated with Ethovision XT (Ver. 07, Noldus, Netherlands). We have conducted one day protocol with a prior habituation of the animal to the behavioral testing room for 30 min and then animals were placed in the central area of the EPM facing either side of the open alley. And then animals were allowed to explore the arena for 5 min. Lastly, the test animals were placed on the central zone facing towards open arm and allowed to explore for five minutes, while time spent and head dipping was recorded. The same was followed in the follow-ups.

### Behavioral scoring with kinoscope for EPM

Apart from the regular analysis of the quantitative variables for elevated plus maze test from the video tracking software (Ethovison XT; Noldus), we also performed post-hoc behavioral scoring from the video tracked by a separate researcher blind to the experiment (intra rater reliability ≥0.9). This is being performed using an available algorithm program ‘kinoscope’ (Kokras et al. 2017) where manually behavioral execution was coded using specific key tagging for different ethograms (viz. exploration to five zones, unprotected head dip), and visual maps were graphically depicted corresponding to individual animals to get an idea about the intra-group variability. These maps of events from start to the end of paradigm (0 to 300 th seconds) were coded with specific color.

### Selection of dose and stereotactic coordinate for injection of STZ

In order to select the dose of STZ, we performed a separate experiment involving three groups with 5 animals/group. AD group animals were injected with three separate doses of STZ, viz. 1.5-, 2- and 3 mg/kg at the lateral ventricles. They were tested for cognitive and anxiety like behavior for 45 days and at the end brain sections were examined by cresyl violet staining. Our observation showed that 3 mg/kg dose is best suited for studying the disease progression for a chronic time period (see Supplementary Fig. S1.b.).

On the other hand, for selection of stereotactic coordinates, we used three separate coordinates and Chicago blue dye was injected using the stereotactic guiding setup (David Kopf, USA), where we found AP: -0.84 mm; ML: ±1.5 mm; DV: -3.5 mm as the best choice as it has good breathing space for surgical manipulation, giving robust coordinates for same precision each time (see supplementary fig. S1.c.).

### Intracerebroventricular injection

In order to induce AD or prepare sham operation, we stereotaxically injected intracerebroventricular STZ (3mg /kg of b/w; bilateral) or aCSF (5µl/site; bilateral) under sodium thiopentone (50 mg/kg of b/w) anesthesia. Injection was performed using a 25 µl Hamilton syringe (22G; 700N glass) placed in arm-held Elite-11 mini pump (Harvard Apparatus, USA) at a rate of 1µl/min, while body temperature was maintained during and after surgery in homeothermic monitoring system (Harvard Apparatus, USA) following previous protocol (Ferry et al. 2014). Briefly, after anesthesia animals were fixed in David-Kopf stereotaxic frame and flat-skull position was achieved cross-checking DV position of Bregma and Lambdoid suture, finally injection was given at AP: -0.84 mm; ML: ±1.5 mm; DV: -3.5 mm coordinates based on rat brain atlas (Paxinos and Charles Watson 2007), through a burr-hole made using a drill-bit (>22G) attached to driller (Foredom, USA). After completion of each injection needle was kept in the same position at least for 3min. Further, postoperative care was taken using meloxicam (Intas pharmaceuticals, India); and gentamicin as analgesic and antibiotic respectively for 3-5 days. aCSF was prepared (Mehla et al. 2012a) and its osmolality checked and maintained at 295±5 mmol/kg to avoid any brain osmotic shock with vapour pressor osmometer (5600; Vapro; Wescor Inc.; USA).

### Transcardial perfusion and tissue harvesting

On the 60^th^ day after terminal behavioral assessments, animals were subjected to 150 mg/kg of sodium thiopentone (i.p.) without any anti-cholinergic drug. After confirmation of complete anesthesia by paw-pressor, animals were placed onto an ice-pack on its back and immediately a lateral incision was made in the skin and abdominal wall and then small incisions were made to access the plural cavity by through incising diaphragm and cardiac envelop was excised to access the heart (Gage et al. 2012). Finally, preceding with ice-cold saline (0.9% NaCl), 50 ml paraformaldehyde (4%) was perfused transcardially. The brain was dissected out and fixed in the same fixative for 3 days at 8°C. The tissues were cryoprotected in 15 to 30% sucrose, 20 µm thick frozen coronal sections taken onto gelatin-coated slides and kept at 8°C until used.

### Histology

The brain sections were stained with cresyl violet (0.4%) for 3 min, dehydrated and mounted with DPX. Congo-red (Sigma Aldrich, USA) staining was performed to assess amyloid plaque (Bott 2014) with counterstaining with haematoxylin. For comparison of hippocampal sections, we restricted to AP-axis from -3.48 to -3.71 mm following the rat brain atlas (Paxinos and Charles Watson 2007). The stained sections were kept in the dark until imaging.

### Golgi-Cox staining

In order to investigate the neurite complexity in the dentate gyrus of the dorsal hippocampus, we performed modified Golgi-Cox staining (Ranjan and Mallick 2010; Zaqout and Kaindl 2016). After terminal behaviour testing, the brain was dissected out and 5 mm chunk of the dorsal hippocampus cut using brain matrix (51388, Stoelting Co., USA) and incubated in Golgi-Cox solution at 37°C for 24 h. They were embedded in 3% agar and 200 µm thick sections cut using vibratome (VT1000S, Leica, GmbH). Sections were mounted onto gelatin-coated slides, developed and finally mounted with DPX.

### Immunostaining

Sections were permeabilized with PBST (0.2% Tween-20 in PBS), blocked in 1% BSA for 2 h before being incubated in anti-GSK3β (1:400; ITT05016, Immunotag, USA), anti-mtCOX (1:200; 1D6E1A8, Abcam, USA), anti-PI3K (1:500; ITT03242, Immunotag, USA) for 2 days at 4°C. Next, sections were incubated with secondary antibodies (anti-rabbit, 1:400; DI-1594; anti-mouse, 1:1000; fl-2000; Vector laboratories, USA) for 2 h. After wash, sections were mounted with fluoroshield with DAPI (F6057, Sigma-Aldrich, USA) and imaged using fluorescence microscopy.

### Image acquisition and analysis

Brain sections stained with cresyl violet, Congo-red and Golgi-Cox were imaged using the brightfield Eclipse Ni upright microscope. For fluorescence imaging, the same microscope was used additionally with fluorescence lamp (C-HGFI, Nikon, Japan) using triple filter block of DAPI (blue; excitation at 385-400 nm; bandpass at 393CWL), FITC (green; excitation at 475-490 nm; bandpass at 483CWL) and TRITC (red; excitation at 545-565 nm; bandpass at 555CWL), according to the need. Images were captured using a colour camera (DSRi-2, Nikon; Japan) and saved in RGB format.

Images of cresyl violet, Golgi-Cox and immunostained sections were analysed for histometric and morphometric evaluations. In case of counting of the number of viable cells and soma areas, we used a specific workflow in the NIS Element basic research software (Nikon, Japan) platform (see supplementary fig.S1.d.). Variables such as lateral ventricular volume, layer wise volumetric measurement, and immunopositive signals were measured/ counted, neurite length, spine density using ‘line tool’ or ‘multi-point tool’ in the FIJI software (Schindelin et al. 2012). All measurements were done in 400X magnification. Cell/ signal counting results were normalized and plotted as ‘per unit area’.

### Statistical analysis

For data distribution, Shapiro-Wilk normality test was performed. For the within group comparison of behaviour with follow-up, one-way ANOVA (non-parametric, two tailed, repeated measures) was performed with Bonferroni’s correction for multiple comparison. For comparison between two groups t-test (nonparametric, Mann-Whitney, two tailed) was performed. For cell count, area of the viable cell and subcellular localization, we performed one-way ANOVA (non-parametric, two tailed, ordinary), in GraphPad Prism 8.0 (GraphPad Software Inc, San Diego, CA). Distribution of swim-time in the four quadrants were analyzed using a continuous multivariate probability distribution named Dirichlet distribution using a python pipeline. The ethograms in EPM test were analyzed and plotted using a specific algorithm (Kinoscope). We performed pairwise correlation of the dataset in case of understanding morpho-functional relationship. A p-value of ≤ 0.05 was considered significant and data were shown as individual values with mean and standard error of mean.

## Results

### Spatial memory impairments after streptozotocin treatment

In the latency to the goal quadrant, we found a significant effect of time (p<0.0001; df= 2; F=20.72), treatment (p<0.0001; df= 4; F= 10.00) as well as the interaction between them (p= 0.0021; df= 8; F= 3.261). Further, on multiple comparisons, latency was found to be increased significantly on 15^th^ (p= 0.0222), 30^th^ (p= 0.0002), 45 (p<0.0001) and 60^th^ day (p= 0.0001) in AD group when compared with its baseline. In the time spent in goal quadrant, two-way ANOVA revealed only significant effect of treatment (p<0.0001; df= 4; F=12.37) and interaction between time and treatment (p= 0.0043; df=8; F= 2.992). Bonferroni’s multiple comparison showed significant reduction of time spent in goal quadrant in sham group on 15^th^ day (p= 0.0027) and 60^th^ day (p= 0.0013) with respect to its baseline, whereas, in AD group, it was reduced on 15^th^ (p= 0.0028), 30^th^ (p<0.0001) 45^th^ (p<0.0001) and 60^th^ day (p<0.0001) when compared with its baseline. In the number of entries to the goal quadrant, only treatment was shown to effect the results significantly (p= 0.0048; df= 4; F= 3.943). Further, on multiple comparison AD group of rats had significantly a smaller number of entries/ visits to the goal quadrant on 30^th^ day (p= 0.0361) and 60^th^ day (p<0.0001) (fig. 1.A-C.).

**Figure 1.:**
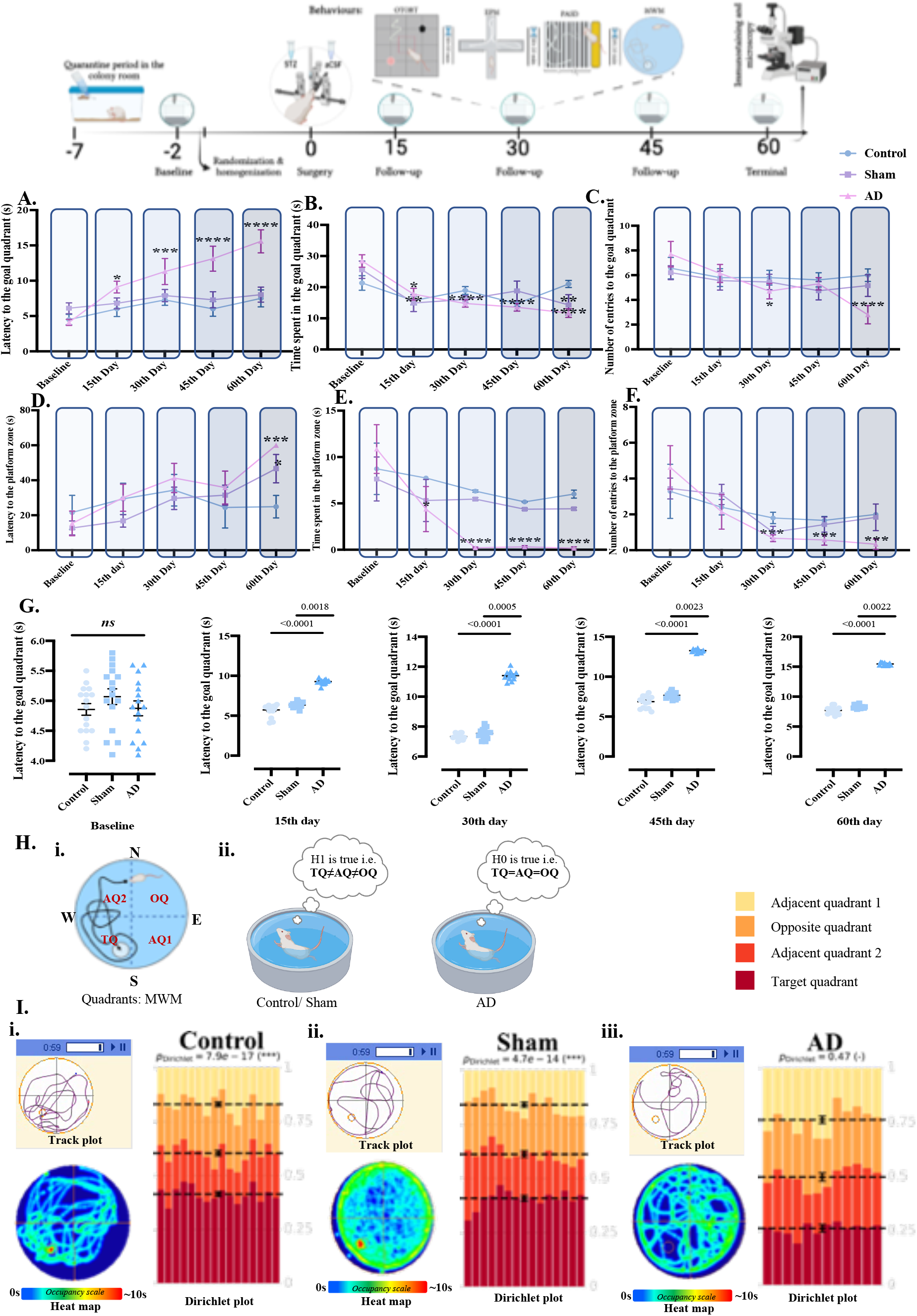
Progressive loss of spatial memory with fraction of time spent in each quadrant does not show an equal or uniform distribution except in AD group of animals as assessed by MWM paradigm. Study design showing the order of behavioural experiments conducted in the study in the top panel; within group analysis of latency to the goal quadrant (A.); time spent in the goal quadrant (B.); number of entries to the goal quadrant (C.); latency to the platform zone (D.); time spent in the platform zone (E.); number of entries in the platform zone (F.); between group comparison of latency to the goal quadrant (G.); template for zone division in MWM tank with the alternative quadrant (AQ), opposite quadrant (OQ) and target quadrant (TQ) made using BioRender application (H.i.); representation of alternative (H1) and null hypothesis (H0) as used in Dirichlet distribution of fraction of time spent in four quadrants in three groups with the colour code used in Dirichlet plot for four quadrants (H.ii); track plot, heat map and Dirichlet plot where each column represents a sample and each colour represents a quadrant. Mean values for the fraction of the time spent in each quadrant is represented by a dotted line and the error bars on the means are approximated with the inverse Fisher information of control (I.i), sham (I.ii) and AD (I.iii.) with n≥ 9; data is expressed as Mean ±SEM except in G, where it is given as Mean ±SEM along with individual data points. * indicates comparison with baseline pre-surgery data ****p≤ 0.0001, *** p≤ 0.001, ** p≤ 0.01, * p≤ 0.05.

On analysis of the latency to the platform zone, we observed only treatment to affect significantly the results (p= 0.0005; df= 4; F= 5.401). Multiple comparison analysis revealed a significant increase in latency to the platform zone on 60^th^ day in sham (p= 0.0211) and AD (p= 0.0007) group when compared with their baseline. In case of time spent in platform we found both time (p= 0.0006; df= 2; F= 7.859) and treatment (p<0.0001; df= 4; F= 8.292) to be significantly different. When multiple comparison was conducted, we observed a decrement on 15^th^ (p= 0.0202), 30^th^ (p<0.0001), 45^th^ (p<0.0001) and 60^th^ day (p<0.0001) in comparison to baseline in AD group only. Number of entries to the platform zone was found to be influenced significantly by treatment (p<0.0001; df= 4; F= 8.660) and multiple comparison test revealed a significant decrement on 30^th^ (p=0.0006), 45^th^ (p= 0.0004) and 60^th^ day (p= 0.0001) in AD group when compared to its baseline (fig. 1.D-F.).

Next, to understand the changes between group at different experimental follow-ups, we performed Kruskal-Wallis test with Dunn’s multiple comparison on latency to the goal quadrant. Statistical analysis revealed no significant mean difference across the groups at baseline pre-surgery data (p= 0.35, df= 2, KW= 2.07). However, Kruskal-Wallis test showed a significant mean difference at 15th (p<0.0001, df= 2, KW= 32.02), 30th (p<0.0001, df= 2, KW= 27.34), 45th (p<0.0001, df= 2, KW= 30.30) and 60th day post-injection of STZ (p<0.0001, df=2, KW= 30.33). Dunn’s multiple comparison on each day revealed a significant increase in AD group as compared to Control (p<0.0001) and Sham (p≤ 0.01; fig. 1.G.). Furthermore, occupancy plot of control and the sham operated groups showed a red or yellowish red in the platform zone respectively, whereas in case of AD group it was found to be blue (fig. 1.I.i-iii.; left panel).

### Distribution of swimming time in the four quadrants of MWM

To probe the strategic changes in the spatial reference memory after i.c.v. STZ injection, we performed likely-ho od ratio statistics (LRS) based on Dirichlet distribution. On 60th day post-injection of STZ, analysis of fraction o f time spent in each quadrant was looked for uniformity within group. Results found a significant uniform distrib ution in Control (p= 7.88896e-17; LRS= 78.0884), Sham (p= 4.74448e-14; LRS= 65.113), except in AD (p= 0.4 73182; LRS= 2.51169) group. Further, as in this statistics we considered equal time spent in each quadrant as nu ll hypothesis (H0) and the alternative hypothesis (H1) was that at least one quadrant will differ significantly. Hen ce, analysis showed that H0 was true only for Control and Sham group of animals (fig. 1.I.i-iii.; right panel).

### Recognition memory deficits after STZ treatment

When we applied repeated measures two-way ANOVA on the recognition index with post hoc Bonferroni’s multiple comparison test, a significant effect of treatment (p<0.0001; df= 4; F= 16.20), time (p<0.0001; df= 2; F= 168.4) and even interaction between the two (p<0.0001; df= 8; F= 13.19) was observed. On multiple comparison between time points, in AD group of rats only, a significant reduction of recognition index on 15^th^ (p<0.0001), 30^th^ (p<0.0001), 45^th^ (p<0.0001), and 60^th^ day (p<0.0001) was evident in comparison to its baseline (fig. 2.A.). In addition, intragroup analysis of discrimination index showed a significant effect of treatment (p<0.0001; df= 4; F= 17.74), time (p<0.0001; df= 2; F= 184.4) and even interaction (p<0.0001; df= 8; F= 14.44). Multiple comparison showed a significant decrease in AD on 15^th^ (p<0.0001), 30^th^ (p<0.0001), 45^th^ (p<0.0001), and 60^th^ day (p<0.0001) as compared to its baseline (fig. 2.B.).

**Figure 2.:**
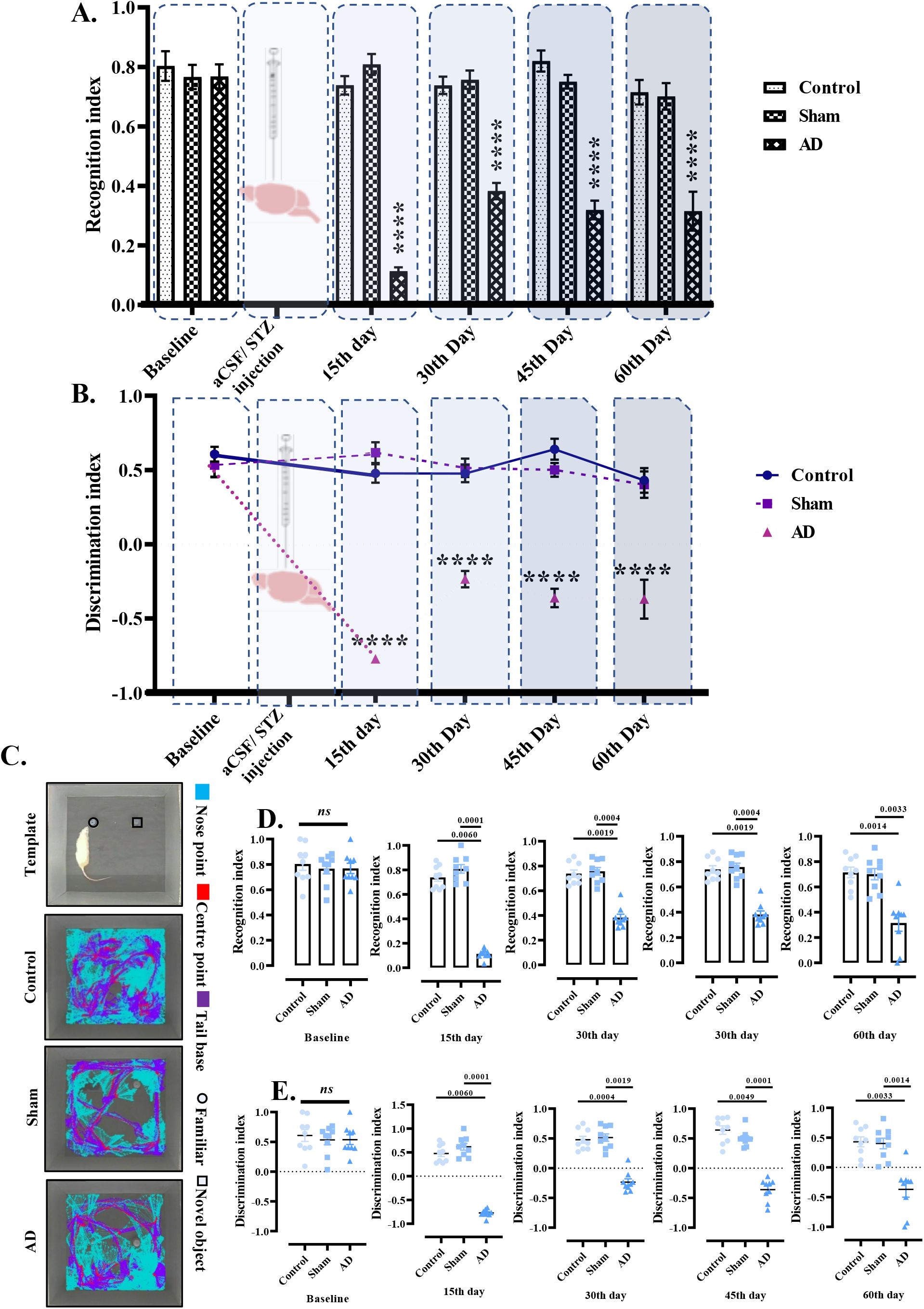
Progressive loss of episodic memory post injection of STZ as assessed by OTORT. within group analysis of recognition index (A.); and discrimination index (B.); track plot as well as template of the original recording frame (C.); between group analysis of recognition index (D.); and discrimination index at various study periods (E.) data is expressed as Mean ±SEM except in D. and E., where it is given as Mean ±SEM along with individual data points; * indicates comparison with baseline pre-surgery data; **** p≤ 0.0001; n≥9/ group.

Furthermore, a significant change in recognition index among the experimental groups examined on 15th (p= 0.0014; df=4; F= 11.87), 30th (p= 0.0002; df= 4; KW= 17.51), 45th (p= 0.0001; df= 4; KW= 18.21) and 60th day post-surgery (p= 0.0005; df= 4; KW= 15.34). No significant difference among the groups was evident at baseline (p= 0.775, df= 4, F= 0.508). Post hoc multiple comparison revealed a significant decrease of recognition index in AD group as compared to the Control (p<0.0001) and sham operated (p≤0.003) groups on all the follow-up days (fig.2.D.). Similarly, between group analysis of discrimination index showed no significant difference among the groups at their baseline (p= 0.775; df= 4; KW= 0.508). However, a significant change was observed on post-surgery day 15 (p= 0.0014; df= 4; F= 11.87), 30 (p= 0.0002; df= 4; KW= 17.51), 45 (p= 0.0001; df= 4; KW=18.21), and 60 (p= 0.0005; df= 4; KW= 15.34). Moreover, multiple comparison revealed significant reduction of discrimination index in AD group as compared to Control (p≤0.001) as well as Sham (p≤0.003) group at all time points (fig.2.E).

Representative track-plots of the of the Control and sham groups on 60th day test phase showed a greater number of line crossing of the nose point (cyan colour), directed/ headed towards novel object in comparison to the familiar one. However, in AD group it was directed towards familiar object, which confirms the results of recognition and discrimination index described in aforementioned paragraph (fig. 2.C.).

### Avoidance memory impairments after STZ treated rats

After training for PASD task at baseline, all rats were tested for retrieval of avoidance memory following STZ injection at an interval of 15 days till 60 days to decipher progression of the disease. Within-group analysis using repeated measures two-way ANOVA with Bonferroni’s multiple comparison test on the latency to the step down revealed a significant influence of interaction between the two (p<0.0001; df= 8; F= 110.1), treatment (p<0.0001; df= 4; F=178.5), and time (p<0.0001; df= 2; F= 1737). Further, post-hoc analysis showed a significant decrease on 45^th^ (p= 0.0169) and 60^th^ day (p= 0.0004) in Sham, and in AD the decrease was significant (p<0.0001) from 15^th^ until 60^th^ day as compared with their baseline (fig. 3.A.).

**Figure 3.:**
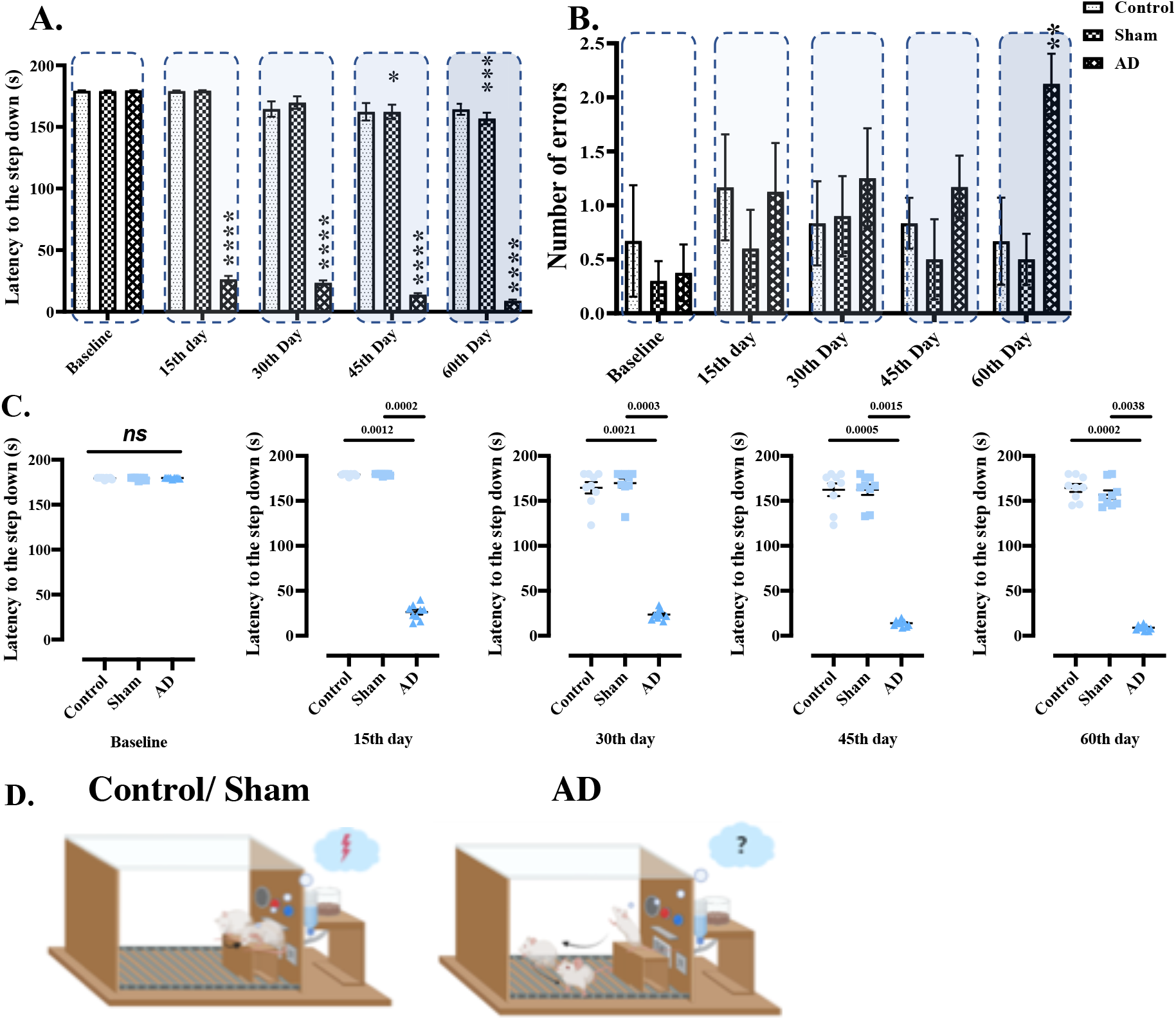
Progressive loss of avoidance memory in STZ-induced AD rats as assessed by passive avoidance step-down paradigm. within group analysis of latency to the step down (A.); and number of errors (B.); illustration of the control/sham group in upper panel and AD group in lower panel created in BioRender app (C.) between group analysis of latency to the step down (D.); data is expressed as Mean ± SEM except in D., where it is given as Mean ±SEM along with individual data points. *indicates comparison with baseline pre-surgery data ****p≤ 0.0001, ***p≤ 0.001, **p≤ 0.01.

Within-group analysis of number of errors at different time-points showed only a significant influence of time (p=0.0220; df=2; F=3.939), and comparison with Tukey’s multiple comparison test showed significant increase in number of errors in AD group on 60^th^ day (p=0.0087) as compared to its baseline (fig. 3.B.).

However, between group analysis failed to show any significant mean differences across the group at baseline (p= 0.75), but post-STZ injection, we found a significant change between the groups examined on 15^th^ (p<0.0001, df= 2; KW= 19.25), 30^th^ (p<0.0001, df= 2, KW= 17.79), 45^th^ (p= 0.0002, df= 2, KW= 17.51) and 60^th^ day (p=0.0001, df=2, KW= 18.00). Further, multiple comparisons revealed significant reduction in AD group versus Control (p≤0.001) and Sham (p≤0.01) group of rats (fig. 3.C.).

### Neuropsychiatric effects seen in STZ-treated rats

On analysis of ratio of time spent in open vs closed arm, we observed time (p= 0.0003; df= 2; F= 8.829) to have a significant influence on the results. However, following multiple comparison, no statistically significant changes with time in any of the groups was observed (fig.4.A.). In head dipping duration, within-group analysis showed a significant influence of treatment (p= 0.0002; df= 4; F= 5.936), time (p<0.0001; df= 2; F= 55.82) as well as interaction between the two factors (p= 0.0096; df=8; F= 2.680). Further, multiple comparison revealed reduction in duration of the head dipping in AD group on 30^th^ (p= 0.0032), 45^th^ (p= 0.0057) and 60^th^ day (p= 0.0001) as compared to its baseline (fig.4.B.).

**Figure 4.:**
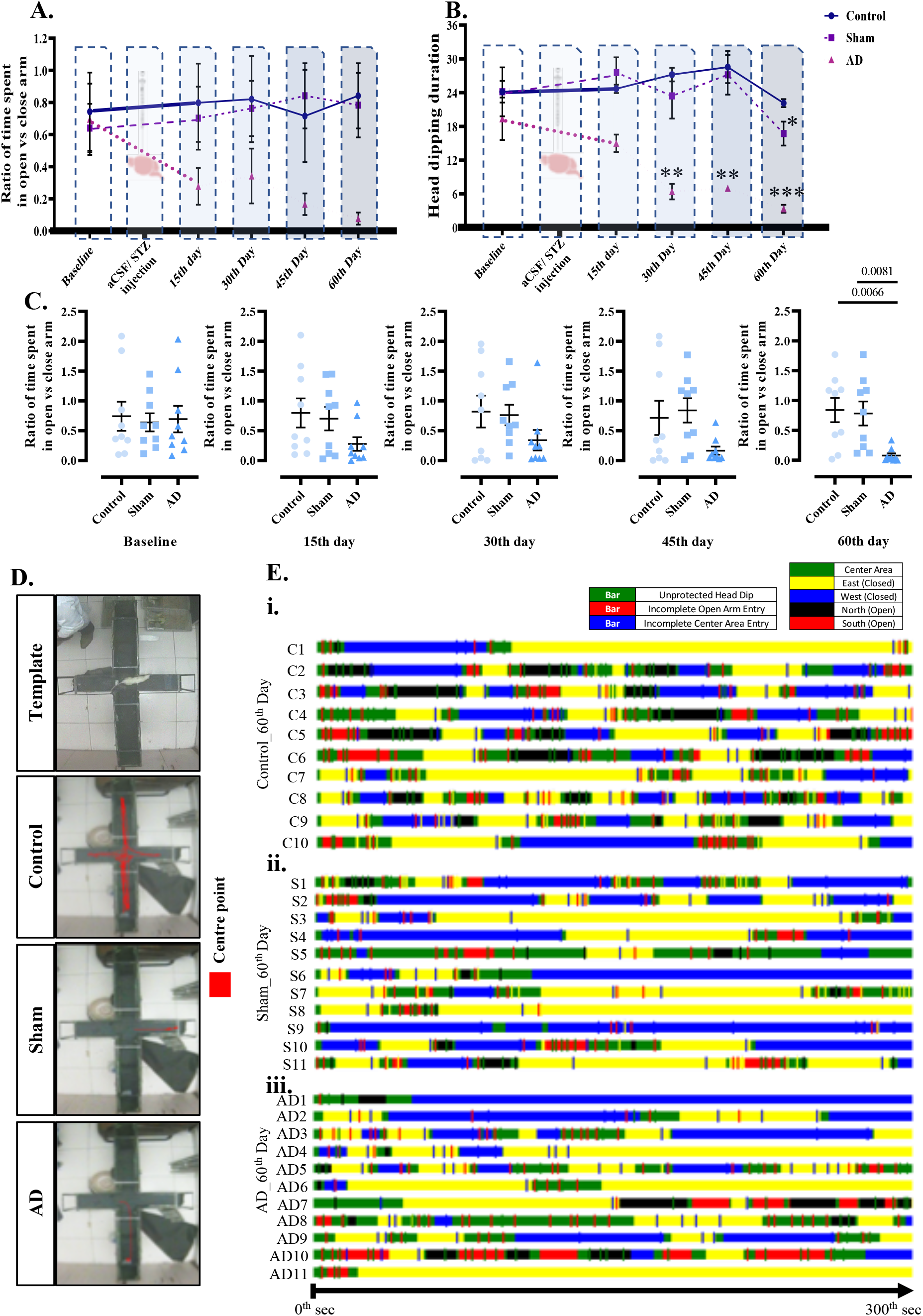
Progressive increase in anxiety after STZ injection with change in the strategy of exploration in EPM. within group analysis of ratio of time spent in open vs closed arm (A.); and head dipping duration (B.); between group ratio of time spent in open vs closed arm (C.); representative track plot of experimental groups with template (D.); ethogram plotted along the time period of terminal behaviour recording on 60^th^ day post-surgery where time spent is represented as column and activity in the arms were represented as bar in control (E.i.), sham (E.ii.), and AD (E.iii.) group; in the post-hoc ethogram analysis with kinoscope software time spent in center area (green), closed (east: yellow; west: blue) arm and open (north: black; south: red) arm are colour coded; data is expressed as Mean ±SEM except in C., where it is given as Mean ±SEM along with individual data points; * indicates comparison with baseline pre-surgery data *** p≤ 0.001, ** p≤ 0.01, * p≤ 0.05.

Kruskal-Wallis test for comparison of data between the groups revealed no significant difference between groups at baseline (p= 0.93, df= 2, KW= 0.13), on 15^th^ (p= 0.095, df= 2, KW= 4.7), 30^th^ day (p= 0.18, df= 2, KW= 3.41) and 45^th^ day (p= 0.12, df= 2, KW= 4.08). Only on 60^th^ day a significant difference was observed (p=0.0022, df= 2, KW= 12.25) across the groups. Further, Dunn’s multiple comparison revealed a significant reduction in the ratio of time spent in open vs closed arm in AD group, when compared with control (p= 0.0066) and sham (p= 0.0081) groups at 60^th^ day follow-up (fig.4.C.).

Moreover, qualitative analysis of anxiety of video track as representative of animal’s exploratory movement in the EPM arena showed that the movement of AD group of rats was restricted more to the closed alley over open alley (fig.4.D.).

### Change in ethograms of STZ-treated animals to anxiety like behaviour

Apart from the within and between group analysis of EPM paradigm, separately we have looked into the strategic changes if taken by the animals at the 60^th^ day post-injection of STZ plotting the visual maps corresponding to the individual animals/ group. Therefore, the post-hoc videos of the terminal day were examined and tagged using kinoscope software where colour coding in the visual map of the Control and Sham group first of all showed an increased number of activities in the form of crossing the five zones as compared to the AD group. Further, in the control, we observed more time spent in the open alley (north or south) labelled with black/red boxes especially in 70% animals (C2-C6, C8-C9), which was accompanied by unprotected head dips (green bars). In case of the sham group, we found fraction of time spent/ exploration in open alley in the 54.5% animals (S1-S2, S5, S7-S9), again accompanied with green bars. Conversely, in case of the AD group of animals, we observed more fraction of time spent in close alley (east or west) labelled as yellow/blue boxes in 72.7% animals (A1-A6, A9, A11) and this behaviour was tightly associated with incomplete open arm entry denoted by red bars (fig.4.E.i-iii.). Therefore, after i.c.v. STZ treatment animals during a trial i.e., 300 second precisely changes its strategic plan to explore the open/ close arm and choses to spend more time in either of the closed arm grooming or prefer to transit from east to west closed arm or vice-versa immediately after an incomplete open arm entry while crossing through the central area or junction of the open and closed arm. Furthermore, 54.5% of animals in the AD group prefer to explore either of the closed arm after placing it on to the central area at the start point. At this initial moment of decisions, we observed stretch-attend posture in majority of the AD induced animals.

### Morphometric changes in the sub-regions of dorsal hippocampus of STZ-injected rat

As hippocampal neurodegeneration leads to cognitive decline, we measured the thickness of the layers juxtaposed to the CA1 sub region of hippocampus in all three experimental groups. We found a layer-wise atrophy in the thickness starting from stratum oriens layer (Or) to the infrapyramidal blade of dentate gyrus (IPBDG) after STZ treatment which has been illustrated superimposed on the cresyl violet stained image (fig.5.A.). Next, we examined third ventricular volume enlargement on cresyl violet stained sections. Where, Kruskal-Wallis statistics revealed a significant change across the group (p=0.0005; df= 2; KW= 11.43) and Dunn’s multiple comparison test exhibited an significant increase in third ventricle of AD group as compared to Control (p=0.014) and Sham (p=0.007) group (fig.5.E.i.). Then, we have measured CA1 layer thickness, and statistical analysis found significant change in left (p= 0.0011; df= 2; KW= 10.78) as well as right (p= 0.0006; df= 2; KW= 11.38) in Kruskal-Wallis test. Dunn’s multiple comparison revealed significant reduction in AD group as compared to Control (right: p= 0.009; left: p= 0.014) and Sham (right: p= 0.011; left: p= 0.012) group. Furthermore, when CA3 layer thickness was examined, we found a significant change in both right (p= 0.0014; df= 2; KW= 10.50) as well as left (p= 0.0004; df= 2; KW= 11.56) hemisphere. In Dunn’s multiple comparison we found a significant loss in thickness of CA3 layer in AD group as compared to Control (right: p= 0.041; left: p= 0.020) Sham (right: p= 0.006; left: p= 0.005) group of animals (fig.5.C.i-iv.). On 60^th^ day post-STZ treatment, two-way ANOVA revealed significant influence of cell type (p<0.0001; df= 2; F=38.98); treatment (p<0.0001; df= 2; F= 243.2) and even interaction between two (p= 0.0039; df=2; F= 6.358) in left hippocampus. Similarly, significant influence of cell type (p<0.0001; df= 1; F= 33.09); treatment (p<0.0001; df= 2; F= 257.2) and even interaction between two (p= 0.0008; df=2; F= 8.475) was observed in right hippocampus. In both right and left hippocampus, Tukey’s multiple comparison revealed a significant reduction of neuronal area both in CA1 as well as in CA3 layer of AD group as compared to control (p<0.0001) and sham (p<0.0001) group (fig.5.C.v-vi.).

**Figure 5.:**
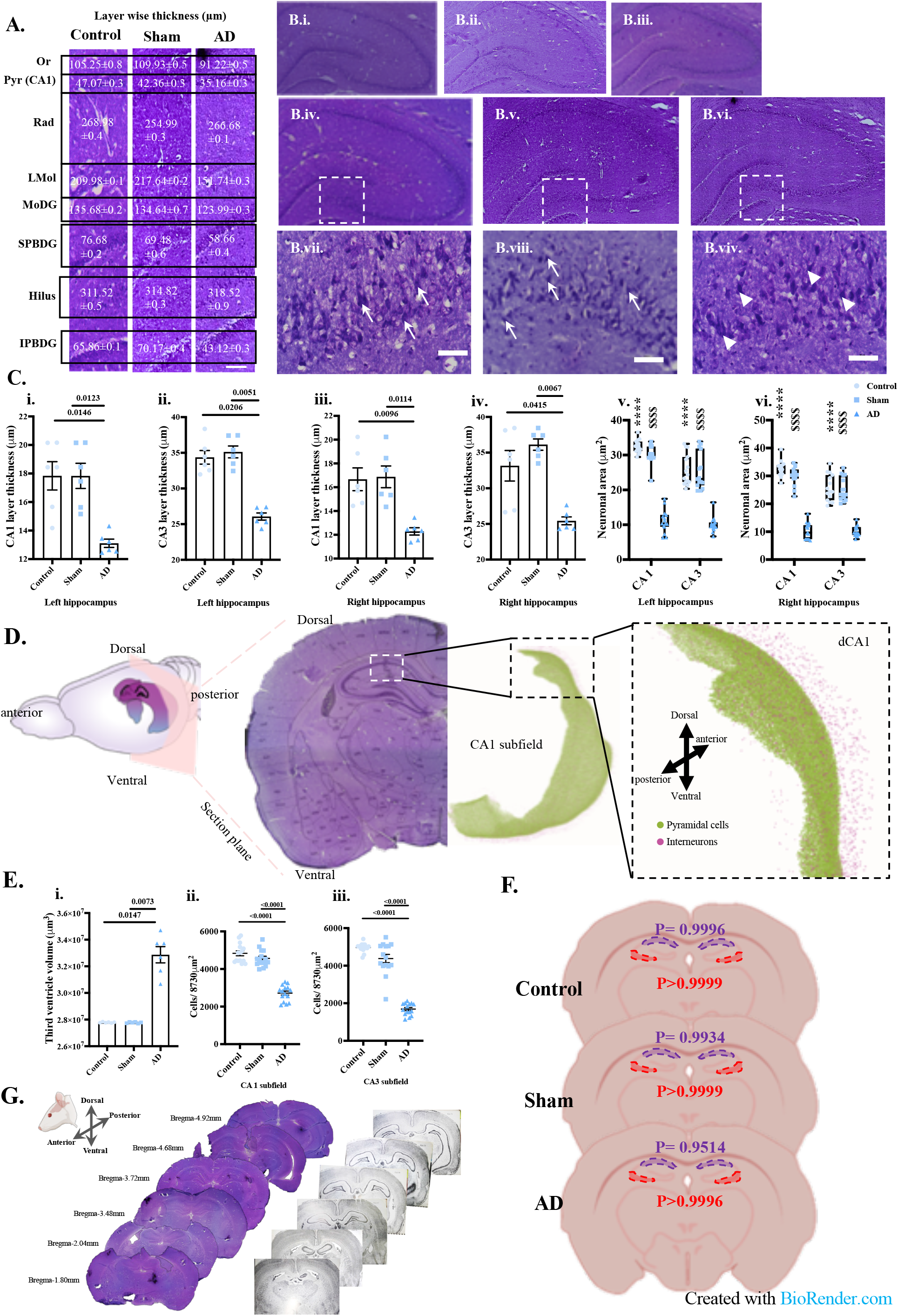
Alteration in the cytoarchitecture of hippocampal sub regions with no lateral biasness and enlargement in cresyl violet stained coronal sections. Layer wise thickness at the vicinity of the CA1 pyramidal layer of hippocampus (scale bar= 500µm; A.); photomicrograph of cresyl violet stained coronal section of Control rat at 4X, 20X and 40X (c.i., c.iv., c.vii.); Sham rat at 4X, 20X and 40X (c.ii., c.v., c.viii.); and AD rat at 4X, 20X and 40X (c.iii., c.vi., c.ix.) showing pycnotic cells with arrow head (scale bar= 50µm; B.); left hippocampal thickness of CA1 layer (C.i.), CA3 layer (C.ii.); right hippocampal thickness of CA1 (C.iii.), and CA3 (C.iv.) layer; neuronal area of left and right hippocampus (C.v-vi.); illustration representing coronal section showing the coronal plane marking the dorsal CA1 representing a detailed neuron model based on Ascoli atlas (https://pair-recording-bsp-epfl.apps.hbp.eu/circuits/rat-ca1) with pyramidal and interneuron (D.) third ventricular volume and cell count in CA1 and CA3 (E.i-iii.); hemispheric-biasness in different experimental groups (F.); selection of anterior-posterior plane to study alterations in AD group (G.); data in neuronal area is expressed as minimum and maximum with individual points whereas, cell count is expressed as Mean±SEM with individual data points; *denotes comparison between control and AD, ^$^denotes comparison between sham and AD group; ^****/$$$$^p≤ 0.0001; n=3 with five fields at 40X magnification (15 data points/ group); or= stratum oriens, Pyr (CA1)= stratum pyramidale of CA1, Rad= stratum radiatum, LMol= lacunusum-moleculare, MoDG= molecular layer of dentate gyrus, SPBDG= suprapyramidal blade of dentate gyrus, IPBDG= infrapyramidal blade of dentate gyrus.

A statistically significant (p<0.0001; df= 2; KW= 30.17) difference among the groups was observed on the cell count of CA1 layer as revealed by Kruskal-Wallis test. Multiple group comparison with Dunn’s test revealed a significant (p<0.0001) reduction in the cell count in AD group as compared to all other group of rats. In case of CA3 layer, cell count also differed significantly (p<0.0001; df= 2; KW= 30.35) among the experimental groups and Dunn’s multiple comparison revealed a significant reduction (p<0.0001) in AD group as compared to other groups (fig.5.E.ii-iii.). All these morphometric data suggest degeneration of CA1 as well as CA3 layer of pyramidal cells in hippocampus of i.c.v. STZ treated rats at 60^th^ day post injection.

Further, we also checked for lateralization or hemispheric-bias if any, for the neuronal area in two hippocampi. Two-way ANOVA analysis showed significant influence of brain areas/ cell types (p<0.0001; df= 3; F= 21.48), treatment/ groups (p<0.0001; df= 2; F= 553.6) and interaction between the two (p=0.0004; df= 6; F= 4.425). However, multiple comparison with Tukey’s test did not show any significant change either in CA1 or in CA3 sub region of hippocampus between the two hemispheric data in any of the experimental groups studied (fig.5.F.).

### Detection of Congophilic particles in the brain of STZ-injected rats

Congo-red stained samples revealed extracellular amyloid plaque deposited in the CA3 sub region of hippocampus of i.c.v. STZ treated rats at 60^th^ day (fig. 6.A.vi.), whereas, in control and sham groups we did not find any such deposits. Further, we have examined other cortical and subcortical areas for the congophilic particles in ICV-STZ injected groups. Upon examination, similar deposits of extracellular particles were present in the primary motor area around layer III/IV and in cingulate gyrus/cortex on 60^th^ day post-injection (fig. 6.B.i). Even in striatal region viz. head of caudate putamen (fig. 6.B.ii.) and corpus callosum and adjacent areas (fig. 6.B.iii.). In addition, these deposits were found in both hemispheres (Fig. 6.C), spanning an area of 3.27 mm in AP axis. These findings underpin that the cognitive impairments observed in behaviour or the neuronal loss in hippocampus in AD group might be tightly related to the deposition of Aβ plaque in various brain areas on 60^th^ day post i.c.v. STZ injection.

**Figure 6.:**
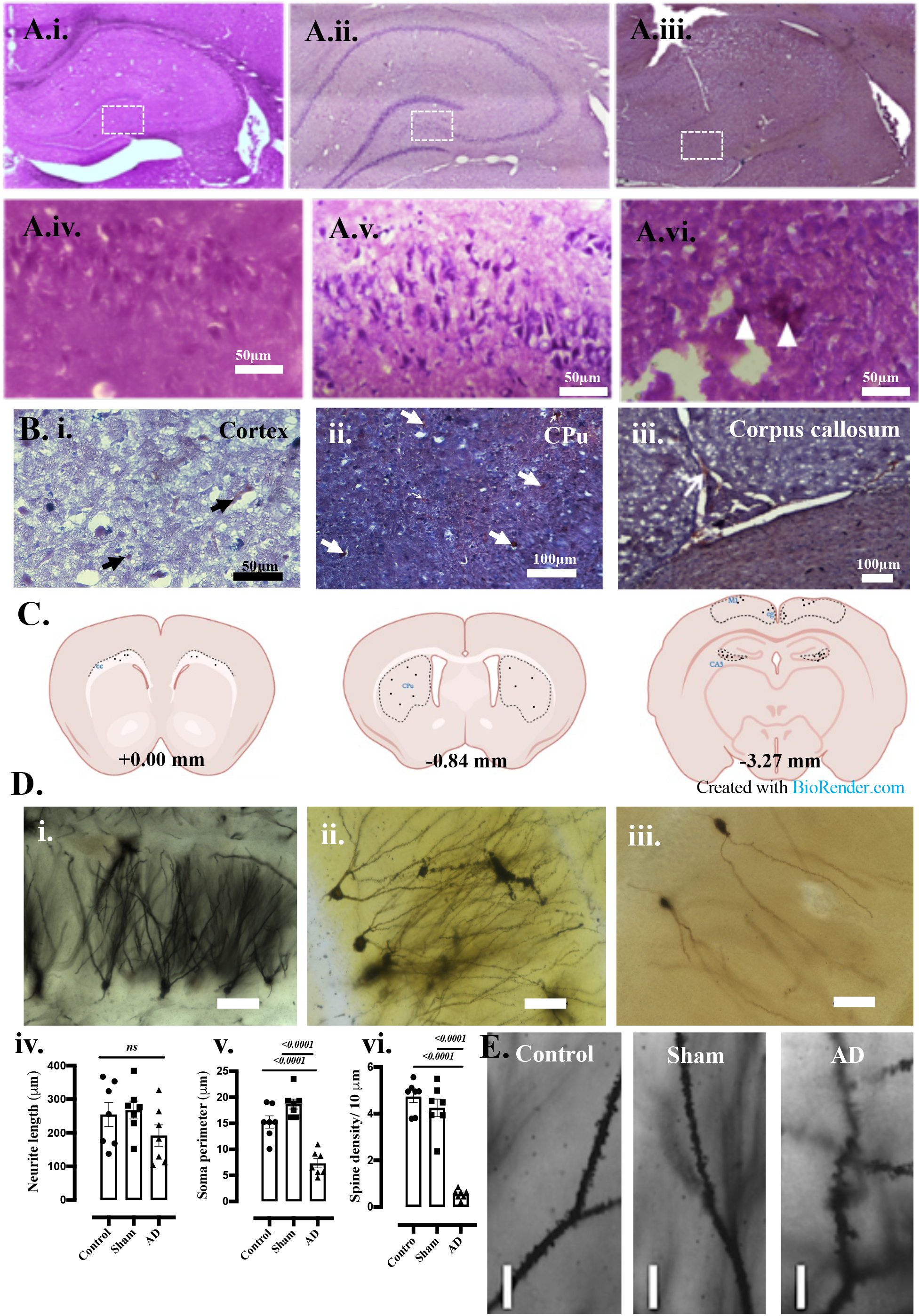
Detection of amyloid plaque in hippocampal formation and impairment in synaptic plasticity in the dentate gyrus of STZ-induced sporadic AD. Photomicrograph of Congo-red stained samples under lower (10X) and higher (40X) magnification of Control (A.i.; A.iv.), Sham (A.ii.; A.v.), and AD (A.iii.; A.vi.) group; Other areas of the brain in AD group showing presence of plaques (B.); schematic illustration of Aβ plaque in various coronal sections of brain of AD group with its AP axis made in BioRender (C.); Golgi-Cox stained granule cells of dendtate gyrus in control (D.i.), sham (D.ii.), and AD (D.iii.); neurite length (D.iv.), soma perimeter (D.v.), spine density (D.vi.) of granule cells; 100X magnification of spines at granule cells of dentate gyrus of control, sham and AD group (scale bar= 7µm; E.); Scale bar= as mentioned on the plates of Congo-red stained samples and 50µm in case of Golgi-Cox stained samples; M1= motor cortex; cg= cingulate gyrus; CPu= caudate putamen; cc= corpus callosum.

### Neuronal and spine morphology of granular cells in STZ-injected rats

In order to understand the synaptic plasticity in the polymorphic layer of hippocampus we measured the neurite length, soma perimeter, spine density of the granular cells after Golgi-Cox staining (fig. 6.D.i-iii.). Ordinary ANOVA test revealed no change (p= 0.225; df= 2; KW= 3.035) in the neurite length across the groups. However, significant changes were observed in the soma perimeter (p<0.0001; df= 2; F= 32.94), where Bonferroni’s multiple comparison revealed a significant reduction in AD group (p<0.0001) as compared to other counterparts. Furthermore, spine density were found to be significantly changed across the group (p<0.0001; df= 2; F= 74.09) and multiple comparison exhibited a significant reduction in the AD group (p<0.0001) as compared to others (fig. 6.D.iv.-vi.; E.).

### Localization of GSK3ß, mtCOX-1 and PI3K in the limbic system of STZ-injected rats

When we compared the adjacent sections of CA1 pyramidal layer of dorsal hippocampus (at -4.36mm) we found a greater number of GSK3β^+^ cells across the whole stratum pyramidale layer (marked with arrow) in AD group. Even in the CA3 layer, we found more expression of GSK3β^+^ cells in AD group as compared to that of Control and Sham counterpart. However, comparison of CA1 and CA3 layer showed elevated expression of GSK3β^+^ cells in CA3 layer with comparison to CA1 layer in AD group. Further, when we looked into the amygdala juxtaposed to piriform cortex and claustrum (at AP -2.28 mm), expression of GSK3β^+^ was much lesser in all sections of Control and Sham group of rats, whereas, the expression was significantly more in the AD group. Similar increase in the GSK3β^+^ cells was also found hypothalamic region juxtaposed to third ventricles in AD group (at AP-2.28 mm), and septum (at AP-0.24 mm) or more specifically bed nucleus of stria terminalis in comparison to Control and Sham group (fig. 7.A: 1.a.-5.c.).

**Figure 7.:**
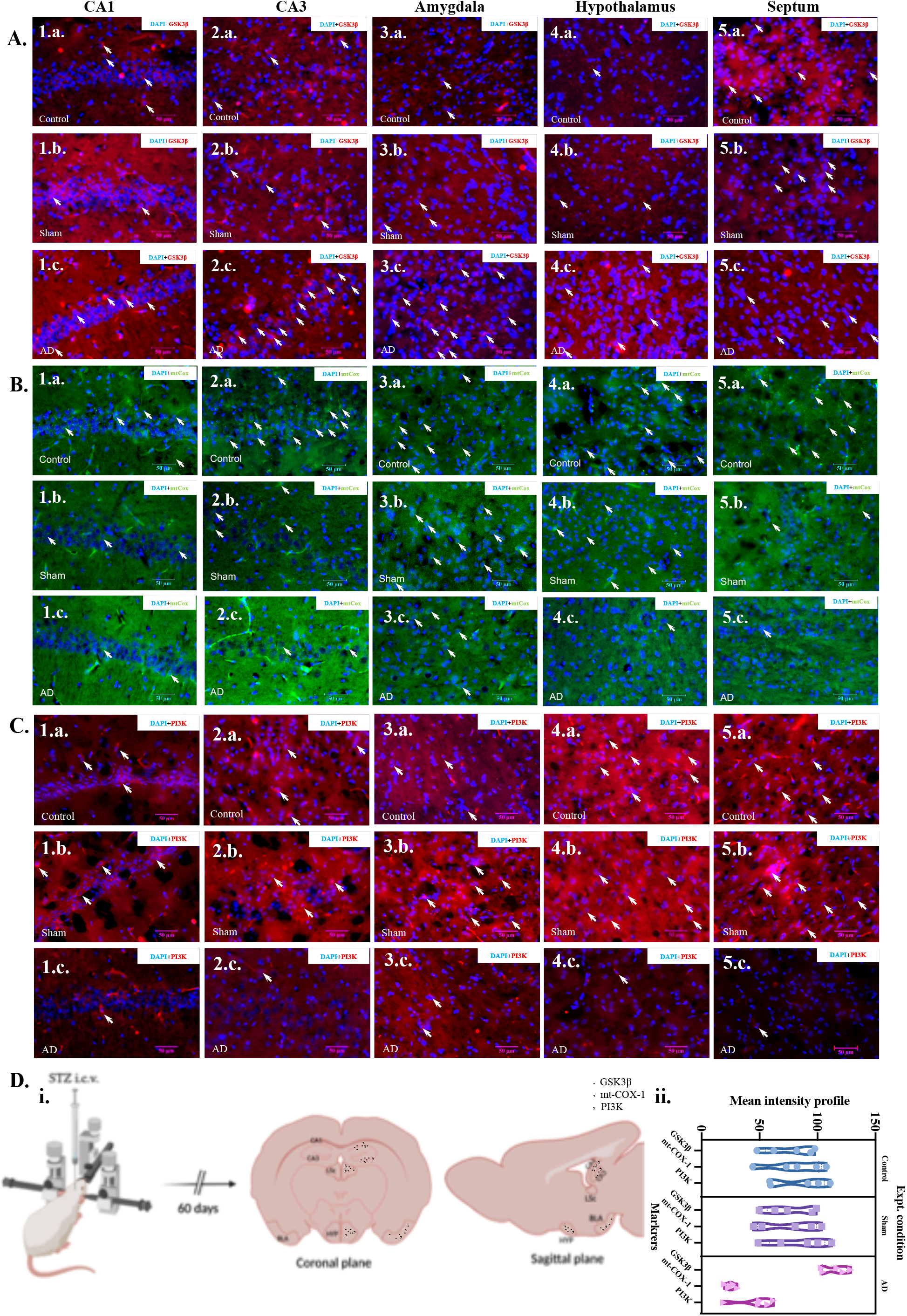
Impairment in the insulin signalling and energy metabolism of brain areas associated with limbic system on two-month post injection of STZ. Photomicrograph of immunostained sections of brain samples on 60^th^ day post-injection of STZ for GSK3β (A.), mt-COX1 (B.), and PI3K (C.) counter stained with DAPI (in blue); 400X magnification images from Control, Sham and AD of CA1 (1.a-1.c.); CA3 (2.a.-2.c.); amygdala (3.a.-3.c.); hypothalamus (4.a.-4.c.); septum (5.a.-5.c.); study design for immunostaining illustrated along with the representative expression of three markers in coronal and sagittal plane of ICV-STZ injected rat (D.i.); mean intensity profile of the three markers are represented as nested plot with experimental groups (D.ii.); Data expressed as violin plot with individual values (n=5/group); CA1= cornu ammonis 1, CA3= cornu ammonis 3, LSc= lateral nucleus of septum, BLA= basolateral nucleus of amygdala, HYP= hypothalamus; scale bar= 50µm.

Further, on comparison, we found an upregulation of mt-COX in the stratum pyramidale and stratum lacunosum layer of CA1 subfield of hippocampus in control as well as Sham group, whereas it was downregulated in the i.c.v. STZ treated rats. Similarly, in CA3 we found a downregulation of mt-COX protein in AD group and in other two groups the expression was more in the stratum oriens and stratum pyramidale. We also observed less expression of mt-COX in amygdala of AD group as compared to Control and Sham group. In the hypothalamus (AP -2.28 mm) and in septum (AP -0.24 mm), we saw a similar upregulation of mt-COX^+^ cells expanding in all layers of the mentioned brain area in Control and Sham group; on the contrary, AD group revealed a downregulation of the same (fig. 7.B; C: 1.a.-5c.). Fig. 7.D.i-ii. Illustrates the expression pattern of three markers and mean intensity profile in nested graph for insulin signalling and mitochondrial activity in five different brain areas after 60^th^ day of STZ treatment.

### Correlation between the morpho-functional variables

Finally in order to understand and predict the exact mechanism of action of STZ for the cognitive and neuropsychiatric changes seen, we performed pairwise correlation between the behavioural and other variables in python. Statistical analysis showed that there is strong correlation between the morphometric and immune parameter studied in the ICV-STZ injected rat brain with their behavioural execution. These correlations proved that findings in the functional (memory and anxiety domain) experiments are tightly associated with the morphological (histometric and subcellular expression of markers) variables on the 60^th^ day post injection, as shown in the correlation heatmap with correlation coefficient values and heat map scale (fig. 8.A.). Further, we performed regression analysis and plotted it between those variables which were found to be more prominently related to each other. There also we have seen the similar association between either morpho-functional or morpho-morpho variables (fig. 8.B.). From these correlation results, we illustrated the predicted mechanism of action of intracerebroventricular injection of STZ in male Wistar rats on 60^th^ day (Fig. 8.C.).

**Figure 8.:**
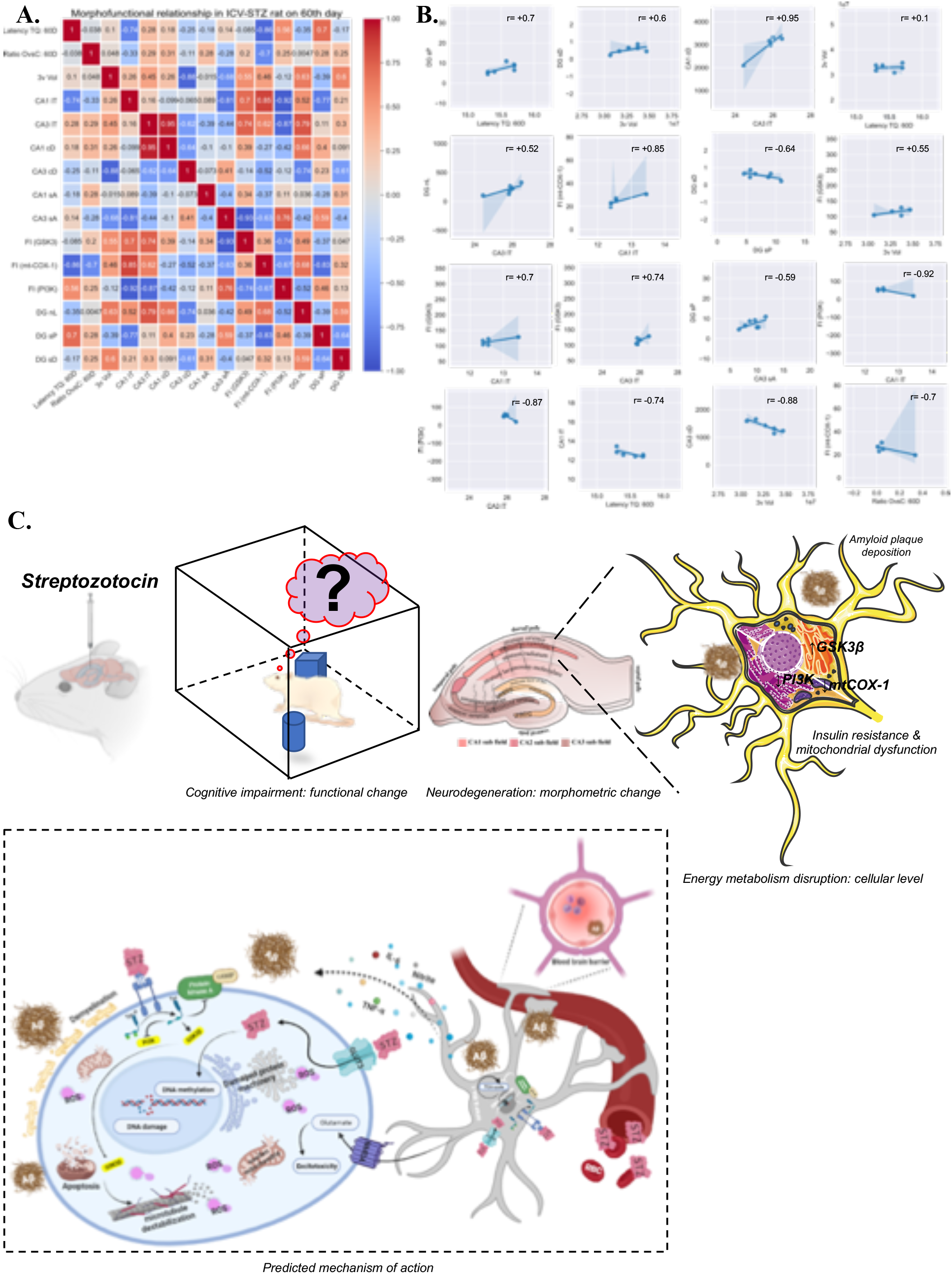
Structure function relationship on 60^th^ day in ICV-STZ injected rats. Correlation matrix with the colour heatmap of ICV-STZ injected rats between functional and morphological variables (A.); reg-plot between important variable found to be strongly related in STZ induced AD rats (B.); proposed mechanism of action of STZ when injected through lateral ventricles (made using BioRender and SMART ppt platform; C.).

## Discussion

There were several reports available on the ICV-STZ injected rats either studied the appropriate dose for phenotypic expression of disease or else studied the effect of novel drugs on the same. On the other hand, the papers talked on the dose-response majorly has selected 3mg/kg of body weight as the best suited dose for progressive cognitive decline. However, work on the pathological phenotyping and its probable reason is elusive. Hence, we have undertaken this research work in order to decipher the reason for the AD like pathology in 3mg/kg b/w (i.c.v.) STZ administered rats.

We have observed a loss of spatial, episodic and avoidance memory in progressive manner from 15th day to the 60th day post-injection of STZ. Further, intergroup comparison revealed this impairment in memory domains are also remain same as compared to that of control and sham operated groups, suggesting a cognitive decline after ICV-STZ injection at 3mg/kg. Further, multivariate analysis of distribution of fraction of time spent in four quadrants of MWM tank on 60th day post-injection of STZ revealed that loss in spatial memory causes changes in swimming strategy, which is similar to that of new learner. This showed that long term memory loss in this animal model is affecting the retention of spatial memory. Bilateral as well as unilateral single or two doses have been shown to induce severe progressive cognitive impairments. Examination of spatial and reference memory in MWM and elevated plus maze for 14 weeks after i.c.v. STZ injection revealed a slow and progressive loss of cognition from two weeks onwards (Mehla et al. 2013; Moreira-Silva et al. 2018a). Cognitive impairment seen in 3mg/Kg i.c.v. STZ injection was associated with overexpression of cytokines at 15th, 18th, 21st day post-injection as assessed with radial arm maze for working and reference memory (Ghosh et al. 2020). In elevated plus maze, a reduction of transfer latency was reported on 17th, 18th, 19th day following twice STZ injection which was correlated with elevated reactive oxygen species and lipid peroxidation (Sharma and Gupta 2001; Kamat et al. 2016).

Unilateral STZ treatment in the left lateral ventricle of WT female mice is shown to have severe loss of episodic memory, which was similar to that of triple transgenic mice model of AD (Chen et al. 2013). In the same study, the recognition memory deficits seen on 14th day post-injection were associated with dysfunction in insulin signaling pathway proteins along with synaptic protein downregulation. In another study by the same group, using unilateral model of STZ injection and 3xTg model, Chen et al. (2012) showed elevation in the expression of 84 genes associated with APP processing, microtubule associated protein tau, synaptic protein, apoptosis/ autophagy, glucose metabolism and mTOR pathway in both the models to a similar extent. These changes were reported to funnel down the cognitive impairment shown after six weeks of STZ injection. Murtishaw et al., (2018) showed that even systemic administration of STZ at 40 mg/kg body wt. on 1, 2, 3, 14, 15 day lead to significant impairment in the discrimination between novel and familiar object exploration, suggesting a neurodegeneration similar to AD and vascular type dementia (Murtishaw et al. 2018). Our OTORT experiments also showed a progressive loss of episodic memory at each follow-up time points in AD group as compared to the Sham and Control group. In a recent study, Moreira-Silva et al. (2018) reported loss of short-term recognition memory on 30th day post injection of STZ, which was associated with loss of synaptic proteins in hippocampus and enlargement of lateral ventricle (Moreira-Silva et al. 2018b). When Wistar rats were tested for object recognition memory after 8th and 14th week post-injection, they showed memory deficit along with upregulation in the phosphor-tau/ tau ratio in the hippocampus (Gáspár et al. 2021). An interesting finding in this paper was that Wistar rats were more prone to have AD like pathological phenotype when compared to the Long-Evans rats, substantiating evidence that selection of right strain for toxin-based animal models of human disease is important. When STZ treated rats were investigated for a stretch of 9 months post-injection for avoidance memory, a slow but steady cognitive deficit in 3mg/kg of body wt. rats was observed exhibiting a biphasic pattern (Knezovic et al. 2015). The pattern of cognitive performance of i.c.v. STZ treated animals showed an immediate severe loss after one week of STZ treatment but an increase on 3rd and decreases from 6-9 month post-injection, the reason was speculated to have an immediate ROS, RNS insult induced by STZ. Though this study focused on long term consequences of STZ injection on cognitive function and pathological staging, they missed out on the time period between 2 weeks-3 months, which according to other studies is the most crucial period wherein severe cognitive as well as morphological deficits are manifested. Further to get a complete picture of the avoidance memory domain in AD, number of errors committed should also have been considered. In our results, we found a severe decline in avoidance memory in PASD test as reflected by step down latency and number of errors at every follow-up time-points starting from 15th day post injection of STZ until 60 days with no biphasic pattern. When single injection of STZ at 3mg/kg of body wt. was administered then a decrease in the avoidance memory in AD group was reported on 14th, 21st and 18th day post injection as compared to control group. This cognitive decline was associated with oxidative stress responses and loss of cholinergic nerve fibers (Mehla et al. 2012b). However, Gáspár et al., (2021) observed avoidance memory at 6th and 11th week post twice (2×1.5mg/kg) or thrice injections (3×1.5mg/kg) of STZ in lateral ventricle and reported no significant change either in step through latency or number of error at any of the follow-up periods (Gáspár et al. 2021). They used shuttle box to study passive avoidance at 6th and 11th week and also gave twice or thrice STZ at a much lower dose of 1.5mg/kg, which may not have been sufficient dosage to induce damage that could result in impairment of avoidance memory.

Next, we looked into the anxiety level in ICV-STZ injected rats, which is seen to be progressively increasing as reflected by decrease in head dipping duration. However, between group comparison showed a significant anxiety only at the 60th day post-injection of ICV-STZ. In the same experiment, ethogram analysis revealed that 72.7% of animals showed an anxious phenotype with spending more time in the close alley of EPM. Again, in this test we observed a change in the strategy of animals induced AD with ICV-STZ avoiding to be inquisitive to suppress the anxiety burden. Even, there were more incomplete transition/ entry in the open arm from the close in these animals suggesting a fearful response making them more conscious of their behavioural execution. Taken together from the quantitative and ethological variables, it is evident that AD group of animals became more conscious/alert while exploring the EPM arena and always have a self-check during transition from one arm to other. This can be best explained by cost-benefit model where animals prefer not to be active/ inquisitive to avoid anxiety burden. Chen et al., (2013) also showed an anxiogenic effect in i.c.v unilateral STZ treated rats on 21st day post-injection, however, they used female mice and earlier experimental evidences has reported effect of sex difference on cognitive function (Chen et al. 2013). Further, no available literature investigated the risk assessment behavior as characterized by head dipping in elevated plus maze. In our results, head dipping duration was found to be reduced with time in AD group, confirming acrophobia induced anxiety in AD. Even a low dose of 1.5mg/kg of body wt. STZ injection in lateral ventricles of mice showed anxiogenic effect in EPM on 7th as well as 21st day post-injection (Pinton et al. 2011). In knock-in model of AD 3xTg female mice enhanced levels of anxiety and fear was observed as assessed by startle responses, freezing behavior and increased restlessness, suggesting an association of neuropsychological behavioral phenotype akin to that seen in human patients (Sterniczuk et al. 2010).

Cognitive impairment in the AD patient or animal models have been directly associated with the neurodegeneration in the dorsal hippocampus (Fuster-Matanzo et al. 2011; Maruszak et al. 2014; O’Callaghan et al. 2019). In present study, we have found appreciable neurodegeneration in ICV-STZ injected group were more restricted to the dorsal hippocampus (fig. 5.E.). This led us to perform histometry of the hippocampus and associated structures viz. layer-wise thickness near to dCA1, morphometry and cell density of CA1 and CA3 layer and third ventricular volume, which showed a neurodegeneration and third ventricular enlargement on 60^th^ day post-injection of STZ. Though inter-hemispheric comparison did not show any lateral biasness. During spatial navigation or exploring novel environment, pyramidal neurons of CA1 and CA3 develop place fields, spatially selective firing fields of place cells, which play an important role in encoding spatial memories. Loss of these neurons leads to spatial memory deficits, as also observed in the present study. In a clinical study where Nissl stained sections were compared for CA1-CA4 of AD brain and age-matched control, prominent neuronal loss was seen only in the CA1 and CA3 pyramidal cell layer of hippocampus, indicating sensitivity of these areas in AD (Padurariu et al. 2012). West et al., (2004) also reported significant neuronal loss in CA1 subfield of hippocampus(48%) as compared other regions; hilus (14%) and subiculum (24%) in AD patients postmortem brain, though in pre-clinical AD cases no significant changes were observed in any of the areas (West et al. 2004). Twice injection (2×1.5mg/kg body wt.) of STZ is reported to have fluoro-Jade C positive cells in CA1 subfield of hippocampus, indicating neuronal degeneration (Yuliani et al. 2020). STZ treatment at a dose of 3mg/kg of body wt. revealed a loss of intact neurons in CA1 pyramidal cell layer on 22^nd^ day post-injection under Cresyl violet staining (Ahmed et al. 2013). Further even gestational systemic administration of STZ at diabetic dose showed a reduction in the density of neurons in CA1 and CA3 but no change in the thickness of the same layers on postnatal 7^th^ and 21^st^ day (Golalipour et al. 2012), confirming neurodegeneration in CA1 and CA3 layers of hippocampus as the hallmark of AD.

A substantial number of reports have demonstrated the presence of amyloid plaques after third month of STZ treatment either by single or twice injection model in various brain areas like hippocampus, cortex, striatum, even in the meningeal capillaries using Congo-red, thioflavin-S and Aβ1-42 antibody staining (Salkovic-Petrisic et al. 2011; Knezovic et al. 2015; Afshar et al. 2018). Hippocampal homogenate of i.c.v. STZ treated rats also showed elevated expression of Aβ1-42 by ELISA or western blot technique from 30^th^ day post-injection (Bao et al. 2017; Shalaby et al. 2019; Ahn et al. 2020; Azadfar et al. 2020). Only in one report reported diffuse amyloid deposition by Congo-red staining on 30^th^ day post injection of STZ though in this study authors did not focused on the hippocampus proper or related structures (Gupta et al. 2018). On examination of coronal sections with Congo-red staining, deposition of congophilic particle at 60^th^ day post-injection was evident not only in CA3 layer of hippocampus but also in cingulate gyrus/cortex, primary motor cortex, caudate putamen and corpus callosum. We did not observe amyloid deposits in capillaries or blood vessels as we speculate that these plaques are first manifested in insulin signaling imbalance prone brain areas and then with disease progression, they traverse through the cerebral meninges showing diffuse and extensive deposition of amyloids more globally, as seen in the long follow-up studies.

Synaptic dysfunction is one of the causes of cognitive decline seen in sporadic AD. The dentate gyrus is the site with neural pluripotent stem cells undergoing adult neurogenesis. Neuroanatomical organization of dentate gyrus revealed an unidirectional afferent fibre towards CA3 layer of hippocampus called mossy fibre tightly associated with episodic memory (Amaral et al. 2007). Experimental body of evidences mainly focused on the CA1 subfield of dorsal hippocampus of human patients (Padurariu et al. 2012). In continuation to the same this subregion of hippocampus also acts as prominent gateway for synaptic inputs to the hippocampal formation (Wosiski-Kuhn and Stranahan 2012). Dendritic spine mainly represent the post-synaptic face of the synapse in-turn transduce changes in the presynaptic neuron in a circuitry (Fischer et al. 1998; Korte and Schmitz 2016). In the present study, we examined the spine morphology of the granular cells and its neurite length, which is found to be damaged on the 60^th^ day of the ICV-STZ injection. This suggests a probable post-synaptic dysfunction in the granule cells of dentate gyrus, site for neurogenesis reflecting a sporadic form of AD. Previous series of papers on STZ induced rat model of diabetes mellites (type I) showed loss of spine in prefrontal cortex, occipital cortex and hippocampus; the loss in hippocampus is not only restricted to the CA1 pyramidal neurons but also to dentate gyrus granular cells and these changes were also extended to dendritic level. These changes were found to be similar to the AD like pathology, which is tightly associated with Aβ1-42 in frontal cortex and hippocampus (Wang et al. 2014). In one recent paper, combination of Golgi-Cox and methoxy-X04 staining in APP/PS1 mouse showed dendritic trimming and spine loss vicinity to amyloid plaques (Kartalou et al. 2020). Another report on 6 month old APP/PS1 mouse model of AD showed that episodic memory deficit in these animals were associated with decrease in LTP with downscaling of spine of mossy fiber from dentate gyrus to CA3 layer of hippocampus (Viana Da Silva et al. 2019). On the other hand, in another article authors showed synaptic dysfunction along with amyloidopathy when STZ is injected into the cisterna magna of rodents at 3mg/kg of body weight (Ahn et al. 2020). In line with that, we observed a strong correlation between the episodic memory variable versus spine density in granular cells of dentate gyrus in ICV-STZ injected rats.

Further, to understand the details of energy metabolism in the STZ treated rats after 60^th^ day post injection we have stained the coronal sections with anti-GSK3β, anti-mt-COX1 antibody and anti-PI3K, which are involved in TCA-cycle or electron transport chain and neuronal survival, respectively. Immunostaining showed an upregulation of the GSK3β and downregulation of mt-COX in the brain areas of limbic system of AD group as compared to Control and Sham group. Unilateral injection of 3mg/kg i.c.v. STZ was found to increase expression of GSK3β in gene as well as protein level at 21^st^ day post-injection in hippocampus and frontal cortex, associated with cognitive and neuropsychiatric dysfunction (Chen et al. 2012, 2013). They also observed imbalance in MAP kinase, PI3K, AKt, presenilin I and BAX, suggesting an insulin signaling dysfunction in the hippocampus as well as frontal cortex. When hippocampal cell line HT-22 was treated with STZ then apart from apoptotic, tau hyperphosphorylation and synaptic loss, Drp1, a mitochondrial fission protein was found to be overexpressed suggesting mitochondrial fragmentation (Park et al. 2020). Even transmission electron microscopic and mitochondrial membrane potential measurement studies showed evidence on alterations in mitochondrial ultrastructure, decrease in transmembrane potential, repolarization level and ATP content after i.c.v. STZ injection (C. Correia et al. 2013; Salkovic-Petrisic et al. 2013; Knezovic et al. 2015).

## Conclusion

Therefore, we conclude from the findings of present study that there is a significant slow and progressive cognitive performances affecting both acquisition and retention of new memory as well as elevate anxiety. These were due to neurodegeneration, amyloid deposition, loss of spine and brain energy imbalance at dose of 3mg/kg at globally involving higher cortex and limbic system. This study not only confirms but also underpins the specific reasons for previous reports on ICV-STZ studies proposing this dose to be the best for long-term investigation. We predict that ICV-STZ animal model is mimicking sporadic form of AD can be a important tool to investigate novel pharmacological and non-pharmacological therapeutic modalities.

## Supporting information

Old supplementary data

New supplementary data

## Abbreviations

aCSF: Artificial cerebrospinal fluid
AD: Alzheimer’s disease
AKT: Protein kinase B
Aß: Amyloid beta
BAX: Bcl-2 associated X protein
DPX: Dibutyl phthalate xylene
Drp1: Dynamin related protein 1
EPM: Elevated plus maze
GSK3ß: Glycogen synthase kinase 3ß
hp-Tau: Hyperphosphorylated tau protein
ICV-STZ: Intracerebroventricular-streptozotocin
IPBDG: Infrapyramidal blade of dentate gyrus
IRBS: Insulin resistance brain state
LMol: Stratum lacunusum-moleculare
MoDG: Molecular layer of dentate gyrus
mt-COX1: Mitochondrial cyclooxygenase 1
mTOR: Mammalian target of rapamycin
MWM: Morris water maze or: Stratum oriens
OTORT: One trial object recognition test
PBST: Tween 20 in phosphate buffered saline
PI3K: Phosphatidyl inositol 3 kinase
PPAR: Peroxisome proliferator activator receptor
Pyr(CA1): Stratum pyramidale of CA1
Rad: Stratum radiatum
SDPA: Stepdown test for avoidance memory
SPBDG: Suprapyramidal blade of dentate gyrus
STZ: Streptozotocin
TQ: Target quadrant

## Author Contributions

Avishek Roy and Suman Jain has conceived the idea and planned the experiment; Avishek Roy written the manuscript with suggestion from Suman Jain, Tapas Chandra Nag; Jantinder Katyal and Yogendra Kumar Gupta supervised cognitive behavioral testing and drug administration, dose selection; Tapas Chandra Nag supervised histological experiment; Sakshi Sharma has conducted Dirichlet distribution and kinoscope analysis with suggestion from Avishek Roy; experiments were carried out and data curated by Avishek Roy and Sakshi Sharma; Avishek Roy performed stereotaxic surgeries and tissue isolation, immunostaining; histometric and other image analysis was performed by Avishek Roy.

## Statement and declaration

### Conflict of interest and ethics approval

Authors in this article claims no potential conflict of interests. Experiments were conducted following guidelines of Institutional Animal Ethical Committee at All India Institute of Medical Sciences, New Delhi vide 937/IAEC/PhD-2016.

### Data availability

Raw data analyzed in the present study will be available with request to lead contact.

## Acknowledgements

We acknowledge Mr Sanjeev Beniwal for assistance in behavioral experiment and Dr Priyanka Kumari, Department of Physiology, All India Institute of Medical Sciences, New Delhi for assistance in histology.

## References

[1.] Afshar S, Shahidi S, Rohani AH, et al (2018) The effect of NAD-299 and TCB-2 on learning and memory, hippocampal BDNF levels and amyloid plaques in Streptozotocin-induced memory deficits in male rats. Psychopharmacology (Berl) 235:2809–2822. https://doi.org/10.1007/s00213-018-4973-x

[2.] Ahmed ME, Moshahid Khan M, Javed H, et al (2013) Amelioration of cognitive impairment and neurodegeneration by catechin hydrate in rat model of streptozotocin-induced experimental dementia of Alzheimerâ€^TM^s type. Neurochem Int 62:492–501. https://doi.org/10.1016/j.neuint.2013.02.006

[3.] Ahn Y, Seo J, Park J, et al (2020) Synaptic loss and amyloid beta alterations in the rodent hippocampus induced by streptozotocin injection into the cisterna magna. Lab Anim Res 36:. https://doi.org/10.1186/s42826-020-00049-x

[4.] Amaral DG, Scharfman HE, Lavenex P (2007) The dentate gyrus: fundamental neuroanatomical organization (dentate gyrus for dummies). Prog Brain Res 163:. https://doi.org/10.1016/S0079-6123(07)63001-5

[5.] Azadfar P, Noormohammadi Z, Noroozian M, et al (2020) Effect of memantine on expression of Bace1-as and Bace1 genes in STZ-induced Alzheimeric rats. Mol Biol Rep 47:5737–5745. https://doi.org/10.1007/S11033-020-05629-7

[6.] Bao J, Mahaman YAR, Liu R, et al (2017) Sex Differences in the Cognitive and Hippocampal Effects of Streptozotocin in an Animal Model of Sporadic AD. Front Aging Neurosci 9:. https://doi.org/10.3389/fnagi.2017.00347

[7.] Bott R (2014) Bancroft’s Theory & Practice of Histological Techniques; 7th ed

[8.] C. Correia S, X. Santos R, S. Santos M, et al (2013) Mitochondrial Abnormalities in a Streptozotocin-Induced Rat Model of Sporadic Alzheimer’s Disease. Curr Alzheimer Res. https://doi.org/10.2174/1567205011310040006

[9.] Chen, Liang Z, Blanchard J, et al (2013) A non-transgenic mouse model (icv-STZ mouse) of Alzheimer’s disease: similarities to and differences from the transgenic model (3xTg-AD mouse). Mol Neurobiol 47:711–725. https://doi.org/10.1007/s12035-012-8375-5

[10.] Chen, Tian Z, Liang Z, et al (2012) Brain Gene Expression of a Sporadic (icv-STZ Mouse) and a Familial Mouse Model (3xTg-AD Mouse) of Alzheimer’s Disease. PLoS One 7:1–12. https://doi.org/10.1371/journal.pone.0051432

[11.] Efird J (2011) Blocked randomization with randomly selected block sizes. Int J Environ Res Public Health 8:15–20. https://doi.org/10.3390/ijerph8010015

[12.] El Sayed NS, Ghoneum MH (2020) Antia, a Natural Antioxidant Product, Attenuates Cognitive Dysfunction in Streptozotocin-Induced Mouse Model of Sporadic Alzheimer’s Disease by Targeting the Amyloidogenic, Inflammatory, Autophagy, and Oxidative Stress Pathways. Oxid Med Cell Longev 2020:. https://doi.org/10.1155/2020/4386562

[13.] Ennaceur A, Cavoy A, Costa JC, Delacour J (1989) A new one-trial test for neurobiological studies of memory in rats. II: Effects of piracetam and pramiracetam. Behav Brain Res. https://doi.org/10.1016/S0166-4328(89)80051-8

[14.] Ennaceur A, Delacour J (1988) A new one-trial test for neurobiological studies of memory in rats. 1: Behavioral data. Behav Brain Res. https://doi.org/10.1016/0166-4328(88)90157-X

[15.] Ferry B, Gervasoni D, Vogt C (2014) Stereotaxic neurosurgery in laboratory rodent: Handbook on best practices

[16.] Filho GLB, Zenki KC, Kalinine E, et al (2015) A new device for step-down inhibitory avoidance task - Effects of low and high frequency in a novel device for passive inhibitory avoidance task that avoids bioimpedance variations. PLoS One. https://doi.org/10.1371/journal.pone.0116000

[17.] Fischer M, Kaech S, Knutti D, Matus A (1998) Rapid actin-based plasticity in dendritic spines. Neuron 20:. https://doi.org/10.1016/S0896-6273(00)80467-5

[18.] Fronza MG, Baldinotti R, Martins MC, et al (2019) Rational design, cognition and neuropathology evaluation of QTC-4-MeOBnE in a streptozotocin-induced mouse model of sporadic Alzheimer’s disease. Sci Rep 9:1–14. https://doi.org/10.1038/s41598-019-43532-9

[19.] Fuster-Matanzo A, Llorens-Martín M, Barreda EG de, et al (2011) Different Susceptibility to Neurodegeneration of Dorsal and Ventral Hippocampal Dentate Gyrus: A Study with Transgenic Mice Overexpressing GSK3β. PLoS One 6:e27262. https://doi.org/10.1371/JOURNAL.PONE.0027262

[20.] Gage GJ, Kipke DR, Shain W (2012) Whole Animal Perfusion Fixation for Rodents. JoVE (Journal Vis Exp e3564. https://doi.org/10.3791/3564

[21.] Gáspár A, Hutka B, Ernyey AJ, et al (2021) Intracerebroventricularly Injected Streptozotocin Exerts Subtle Effects on the Cognitive Performance of Long-Evans Rats. Front Pharmacol 0:1130. https://doi.org/10.3389/FPHAR.2021.662173

[22.] Ghosh R, Sil S, Gupta P, Ghosh T (2020) Optimization of intracerebroventricular streptozotocin dose for the induction of neuroinflammation and memory impairments in rats. Metab Brain Dis 35:1279–1286. https://doi.org/10.1007/s11011-020-00588-1

[23.] Golalipour MJ, Kafshgiri SK, Ghafari S (2012) Gestational diabetes induced neuronal loss in CA1 and CA3 subfields of rat hippocampus in early postnatal life. Folia Morphol 71:

[24.] Grünblatt E, Salkovic-Petrisic M, Osmanovic J, et al (2007) Brain insulin system dysfunction in streptozotocin intracerebroventricularly treated rats generates hyperphosphorylated tau protein. J Neurochem 101:757–770. https://doi.org/10.1111/j.1471-4159.2006.04368.x

[25.] Gupta S, Yadav K, Mantri SS, et al (2018) Evidence for Compromised Insulin Signaling and Neuronal Vulnerability in Experimental Model of Sporadic Alzheimer’s Disease. Mol Neurobiol 55:8916–8935. https://doi.org/10.1007/S12035-018-0985-0

[26.] Homolak J, Kodvanj I, Babic Perhoc A, et al (2022) Nitrocellulose redox permanganometry: A simple method for reductive capacity assessment. MethodsX 9:. https://doi.org/10.1016/J.MEX.2021.101611

[27.] Kamat PK, Kalani A, Rai S, et al (2016) Streptozotocin Intracerebroventricular-Induced Neurotoxicity and Brain Insulin Resistance: a Therapeutic Intervention for Treatment of Sporadic Alzheimer’s Disease (sAD)-Like Pathology. Mol. Neurobiol. 53:4548–4562

[28.] Kartalou GI, Endres T, Lessmann V, Gottmann K (2020) Golgi-Cox impregnation combined with fluorescence staining of amyloid plaques reveals local spine loss in an Alzheimer mouse model. J Neurosci Methods 341:. https://doi.org/10.1016/J.JNEUMETH.2020.108797

[29.] Knezovic A, Osmanovic-Barilar J, Curlin M, et al (2015) Staging of cognitive deficits and neuropathological and ultrastructural changes in streptozotocin-induced rat model of Alzheimer’s disease. J Neural Transm 122:577–592. https://doi.org/10.1007/s00702-015-1394-4

[30.] Kokras N, Baltas D, Theocharis F, Dalla C (2017) Kinoscope: An open-source computer program for behavioral pharmacologists. Front Behav Neurosci 11:88. https://doi.org/10.3389/FNBEH.2017.00088/BIBTEX

[31.] Korte M, Schmitz D (2016) CELLULAR AND SYSTEM BIOLOGY OF MEMORY : TIMING, MOLECULES, AND BEYOND CELLULAR BIOLOGY OF MEMORY SYSTEMS BIOLOGY OF MEMORY. Physiol Rev 96:647–693. https://doi.org/10.1152/physrev.00010.2015

[32.] Lin LF, Jhao YT, Chiu CH, et al (2022) Bezafibrate Exerts Neuroprotective Effects in a Rat Model of Sporadic Alzheimer’s Disease. Pharmaceuticals 15:. https://doi.org/10.3390/ph15020109

[33.] Maruszak A, Thuret S, Mueller SG, Boccia MM (2014) Why looking at the whole hippocampus is not enough-a critical role for anteroposterior axis, subfield and activation analyses to enhance predictive value of hippocampal changes for Alzheimer’s disease diagnosis. Front Cell Neurosci 8:. https://doi.org/10.3389/fncel.2014.00095

[34.] Maugard M, Doux C, Bonvento G, Morris RGM (2019) new statistical method to analyze Morris Water Maze data using Dirichlet distribution [version 2 ; peer review : 2 approved]. F1000Research 8:1–14

[35.] Mehla J, Pahuja M, Dethe SM, et al (2012a) Amelioration of intracerebroventricular streptozotocin induced cognitive impairment by Evolvulus alsinoides in rats: In vitro and in vivo evidence. Neurochem Int 61:1052–1064. https://doi.org/10.1016/J.NEUINT.2012.07.022

[36.] Mehla J, Pahuja M, Dethe SM, et al (2012b) Amelioration of intracerebroventricular streptozotocin induced cognitive impairment by Evolvulus alsinoides in rats: In vitro and in vivo evidence. Neurochem Int 61:1052–1064. https://doi.org/10.1016/J.NEUINT.2012.07.022

[37.] Mehla J, Pahuja M, Gupta YK (2013) Streptozotocin-induced sporadic Alzheimer’s Disease: Selection of appropriate dose. J Alzheimer’s Dis. https://doi.org/10.3233/JAD-2012-120958

[38.] Moreira-Silva D, Carrettiero DC, Oliveira ASA, et al (2018a) Anandamide Effects in a Streptozotocin-Induced Alzheimer’s Disease-Like Sporadic Dementia in Rats. Front Neurosci 0:653. https://doi.org/10.3389/FNINS.2018.00653

[39.] Moreira-Silva D, Carrettiero DC, Oliveira ASA, et al (2018b) Anandamide Effects in a Streptozotocin-Induced Alzheimer’s Disease-Like Sporadic Dementia in Rats. Front Neurosci 0:653. https://doi.org/10.3389/FNINS.2018.00653

[40.] Morris R (1984) Developments of a water-maze procedure for studying spatial learning in the rat. J Neurosci Methods. https://doi.org/10.1016/0165-0270(84)90007-4

[41.] Murtishaw A, Heaney C, Bolton M, et al (2018) Intermittent streptozotocin administration induces behavioral and pathological features relevant to Alzheimer’s disease and vascular dementia. Neuropharmacology 137:164–177. https://doi.org/10.1016/J.NEUROPHARM.2018.04.021

[42.] O’Callaghan C, Shine JM, Hodges JR, et al (2019) Hippocampal atrophy and intrinsic brain network dysfunction relate to alterations in mind wandering in neurodegeneration. Proc Natl Acad Sci 116:3316–3321. https://doi.org/10.1073/PNAS.1818523116

[43.] Padurariu M, Ciobica A, Mavroudis I, et al (2012) HIPPOCAMPAL NEURONAL LOSS IN THE CA1 AND CA3 AREAS OF ALZHEIMER’S DISEASE PATIENTS. Psychiatr Danub 24:152–158

[44.] Park J, Won J, Seo J, et al (2020) Streptozotocin Induces Alzheimer’s Disease-Like Pathology in Hippocampal Neuronal Cells via CDK5/Drp1-Mediated Mitochondrial Fragmentation. Front Cell Neurosci 14:. https://doi.org/10.3389/fncel.2020.00235

[45.] Paxinos G, Charles Watson (2007) The Rat Brain in Stereotaxic Coordinates Sixth Edition

[46.] Pinton S, da Rocha J, Gai B, Nogueira C (2011) Sporadic dementia of Alzheimer’s type induced by streptozotocin promotes anxiogenic behavior in mice. Behav Brain Res 223:1–6. https://doi.org/10.1016/J.BBR.2011.04.014

[47.] Ranjan A, Mallick BN (2010) A modified method for consistent and reliable Golgi-Cox staining in significantly reduced time. Front Neurol. https://doi.org/10.3389/fneur.2010.00157

[48.] Rostami F, Javan M, Moghimi A, et al (2017) Streptozotocin-induced hippocampal astrogliosis and insulin signaling malfunction as experimental scales for subclinical sporadic Alzheimer model. Life Sci 188:172–185. https://doi.org/10.1016/j.lfs.2017.08.025

[49.] Salkovic-Petrisic M, Knezovic A, Hoyer S, Riederer P (2013) What have we learned from the streptozotocin-induced animal model of sporadic Alzheimer’s disease, about the therapeutic strategies in Alzheimer’s research. J Neural Transm 120:233–252. https://doi.org/10.1007/s00702-012-0877-9

[50.] Salkovic-Petrisic M, Osmanovic-Barilar J, Brückner MK, et al (2011) Cerebral amyloid angiopathy in streptozotocin rat model of sporadic Alzheimer’s disease: A long-term follow up study. J Neural Transm 118:765–772. https://doi.org/10.1007/s00702-011-0651-4

[51.] Salkovic-Petrisic M, Tribl F, Schmidt M, et al (2006) Alzheimer-like changes in protein kinase B and glycogen synthase kinase-3 in rat frontal cortex and hippocampus after damage to the insulin signalling pathway. J Neurochem 96:1005–1015. https://doi.org/10.1111/j.1471-4159.2005.03637.x

[52.] Schindelin J, Arganda-Carreras I, Frise E, et al (2012) Fiji: An open-source platform for biological-image analysis. Nat. Methods

[53.] Shalaby MA, Nounou HA, Deif MM (2019) The potential value of capsaicin in modulating cognitive functions in a rat model of streptozotocin-induced Alzheimer’s disease. Egypt J Neurol Psychiatry Neurosurg 2019 551 55:1–13. https://doi.org/10.1186/S41983-019-0094-7

[54.] Sharma, Gupta Y (2001) Intracerebroventricular injection of streptozotocin in rats produces both oxidative stress in the brain and cognitive impairment. Life Sci 68:1021–1029. https://doi.org/10.1016/S0024-3205(00)01005-5

[55.] Sterniczuk R, Antle MC, Laferla FM, Dyck RH (2010) Characterization of the 3xTg-AD mouse model of Alzheimer’s disease: Part 2. Behavioral and cognitive changes. Brain Res 1348:149–155. https://doi.org/10.1016/J.BRAINRES.2010.06.011

[56.] Viana Da Silva S, Zhang P, Haberl MG, et al (2019) Hippocampal mossy fibers synapses in CA3 pyramidal cells are altered at an early stage in a mouse model of Alzheimer’s disease. J Neurosci 39:4193–4205. https://doi.org/10.1523/JNEUROSCI.2868-18.2019

[57.] Walf AA, Frye CA (2007) The use of the elevated plus maze as an assay of anxiety-related behavior in rodents. Nat Protoc. https://doi.org/10.1038/nprot.2007.44

[58.] Wang JQ, Yin J, Song YF, et al (2014) Brain aging and AD-like pathology in streptozotocin-induced diabetic rats. J Diabetes Res 2014:. https://doi.org/10.1155/2014/796840

[59.] West MJ, Kawas CH, Stewart WF, et al (2004) Hippocampal neurons in pre-clinical Alzheimer’s disease. Neurobiol Aging 25:1205–1212. https://doi.org/10.1016/J.NEUROBIOLAGING.2003.12.005

[60.] Wosiski-Kuhn M, Stranahan AM (2012) Transient increases in dendritic spine density contribute to dentate gyrus long-term potentiation. Synapse 66:661–664. https://doi.org/10.1002/syn.21545

[61.] Xu Z-P, Li L, Bao J, et al (2014) Magnesium Protects Cognitive Functions and Synaptic Plasticity in Streptozotocin-Induced Sporadic Alzheimer’s Model. PLoS One 9:e108645. https://doi.org/10.1371/journal.pone.0108645

[62.] Yuliani T, Lobentanzer S, Klein J (2020) Central cholinergic function and metabolic changes in streptozotocin-induced rat brain injury. J Neurochem. https://doi.org/10.1111/JNC.15155

[63.] Zaqout S, Kaindl AM (2016) Golgi-cox staining step by step. Front Neuroanat. https://doi.org/10.3389/fnana.2016.00038

