## Supplementary material for "Cognitive dysfunction and anxiety resulting from synaptic downscaling, hippocampal atrophy and ventricular enlargement with intracerebroventricular streptozotocin injection in male Wistar rats": Old supplementary data

### Supplementary document

1. **Selection of objects for the one trial object recognition test (OTORT):**
In order to select object for the OTORT we first set out to find out the interest index of 6 control animals for eight different objects (fig. S1.a.A.), out of which one has been selected for the present study to use as novel object. Interest index is calculated by dividing the time spent in exploring novel object with total time allowed to explore the arena i.e. 300s. For the same, we have first habituated the animals with the arena and then we have first went for a sample phase and test phase. The protocol is same as described in the method section (Ennaceur et al., 1989) of the main text (fig. S1.a.C). The representative track plot of sample phase (fig. S1.a.B.i) and test phase of 8 objects (fig. S1.a.B.ii-ix). Furthermore, exploration time for familiar and novel object and interest index of 8 objects tested for novelty was plotted as histogram in fig. S1.a.D-E.

2. **Selection of the dose of streptozotocin (STZ):**
In order to select the appropriate dose of STZ in the present study we have studied effect of three doses (1.5; 2.5; and 3mgg/kg of bw.) and injected using the same protocol as described in the method section in main text (Ferry et al., 2014). Further, we have conducted Morris water maze test for 45 days post-injection of STZ and tried to understand the dose response where statistical analysis showed two major things (n=5/group). Firstly, inter group comparison at different time points showed a significant increase in the latency to the platform zone in all three ICV-STZ injected group as compared to the sham injected rats ($p<0.0001$; fig. S1.b.i). Secondly, comparison between only three STZ injected groups exhibited significant decrease in 30$^{th}$ (vs. 15$^{th}$ day: p= 0.038) and 45$^{th}$ day (vs. 15$^{th}$ day: p= 0.002; vs. 30$^{th}$ day: p= 0.0005) in 1.5mg/kg group, whereas increase only on 45$^{th}$ day (vs. 30$^{th}$ day: p= 0.011) was seen in 2.5mg/kg group. On the other hand, in 3mg/kg ICV-STZ injected group we found an increase on 30$^{th}$ day (vs. 15$^{th}$ day: p= 0.0047) as well as on 45$^{th}$ day (vs. 15$^{th}$ day: p= 0.0003; vs. 30$^{th}$ day: $p<0.0001$) as exhibited by Bonferroni's multiple comparison. In this experiment a progressive increase in response with steep slope of the curve was noted as shown in the spaghetti-line graph in figure S1.b.ii. At the end of the 45$^{th}$ day blood was collected and serum was isolated for nitrocellulose redox permanganometry (NPR) as mentioned earlier (Homolak et al., 2022) of serum samples (n=2/group; 6 biological replicates) showed a significant increase in oxidative stress in the serum of the 3mg/kg ICV-STZ as compared to 1.5mg/kg (p= 0.042) and sham ($p<0.0001$) injected group on 45$^{th}$ day post-injection of ICV-STZ. However, 2.5mg/kg UCV-STZ injected group also showed increase in NPR as compared to sham (p= 0.042) group (fig. S1.b.iii.). And cresyl violet staining on the other hand in the remaining animals (n=3/group) showed a pycnosis in all the ICV-STZ injected rats CA3 subfield of hippocampus as indicated by arrow heads, however the number of the viable cells were least in the 3mg/kh group (fig. S1.b.iv.a-d). Taken together these finding suggests that 3mg/kg ICV-STZ injection is the best choice of drug dose to study the disease progression for two month period.

3. **Selection of stereotactic coordinate for injection of streptozotocin:**
In order to select the coordinate for stereotaxic surgery to inject STZ intracerebroventricularly, we have injected Chicago blue dye (Central drug house, India) using the stereotaxic injection protocol as described in the method section (Ferry et al., 2014) for stereotaxic injection (fig.S1.c.i) in three stereotaxic coordinates viz. AP-0.12mm, ML±1.5mm, DV-3.5mm; AP-0.84mm, ML±1.5mm, DV-3.5mm; AP-1.80mm, ML±1.5mm, DV-3.5mm (fig. S1.c.iii.) in three separate animals. We have reached exactly in the lateral ventricle only in the second AP

coordinate as mentioned earlier, which is shown in images of the 1mm chunks of coronal sections of fixed brains sectioned in brain matrix (Stoelting Co., USA; fig. S1.c.1.b).

4. **Workflow for counting of viable cells:**
we have counted viable cells as well as the soma area using the thresholding option in the NIS basic research analysis (NIKON Instruments, Japan) program available with the NIKON Eclipse microscope in the coronal sections stained with cresyl violet. First images were taken at 40X objective (400X magnification) for CA1, CA2, CA3 subfield of hippocampus. In brief, first image was opened in the software, and then with the Bezier drawing tool CA1, CA2 and CA3 were selected manually as region of interest (ROI). Next, ROIs were thresholded with color threshold plugin and noises were deselected carefully. For segmentation of the noise and signals we have used four separate digital filters available in the NIS Element BR software namely Smooth, Clean, Fill holes, Separate. Finally, measurement was performed for the field and data was exported to '.xlx' format. Which was further compiled for analysis and statistical tests which has been stepwise shown in figure S1.d.

**Figure legends:**

## S1.a.

Formulae for interest index (II): II=($T_N$/$T_{AE}$) Where, $T_N$= Time spent in exploring novel object; $T_{AE}$= Time allowed to explore the arena i.e. 300sec

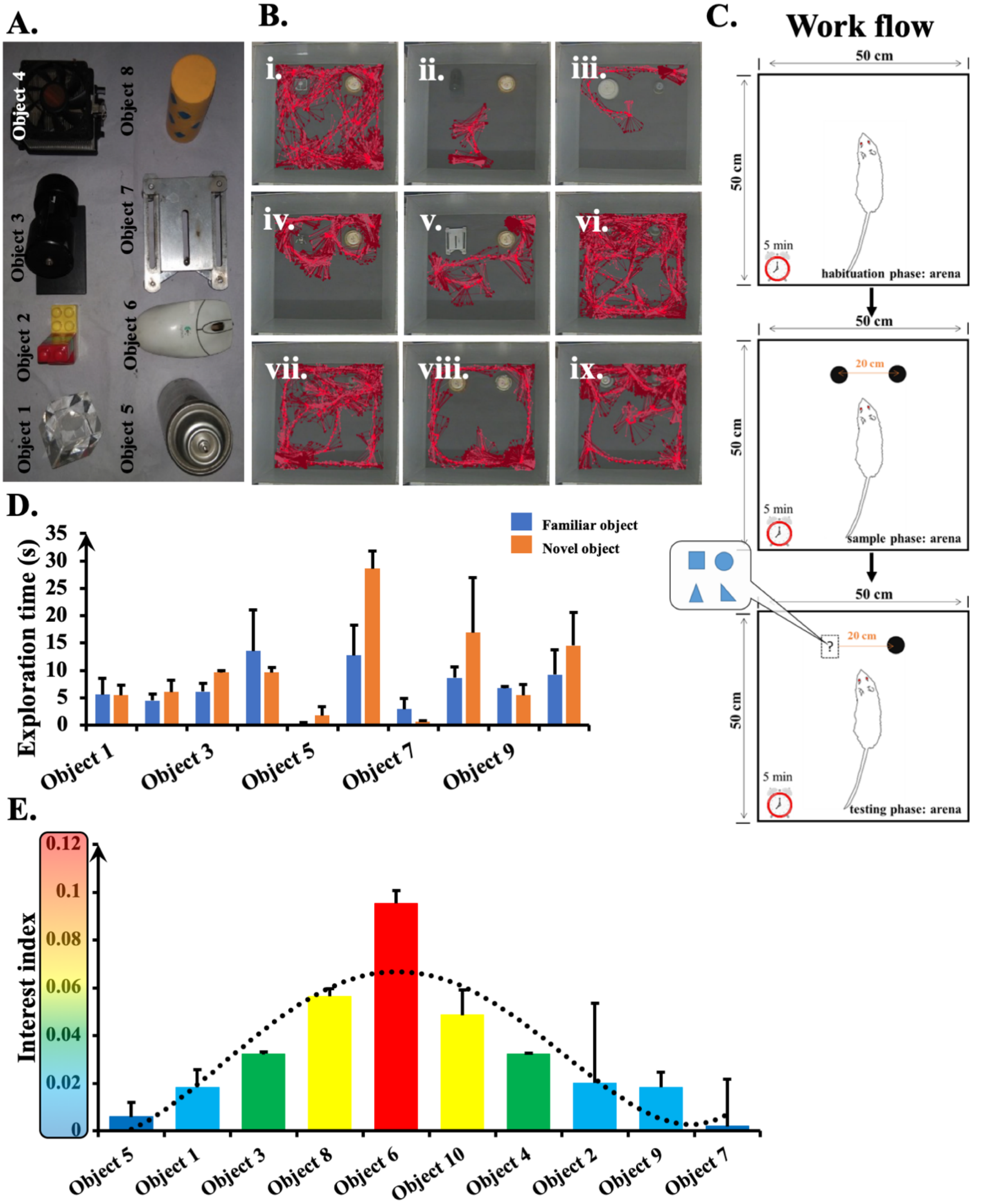


**Figure S1.a.: Selection of novel object for the one trial object recognition test.**
Different objects used in the experiment to select novel object (A); track plot of the sample phase and different testing phase of the eight objects used for novel object selection procedure (B.i-ix.); workflow of the selection of novel object step-wise (C); exploration time exploring

the familial and novel object in the experiment against each object (D); interest index calculated from the formulae mentioned at the top of the figure (E); data expressed as mean±SEM (n=6).

**S1.b.**

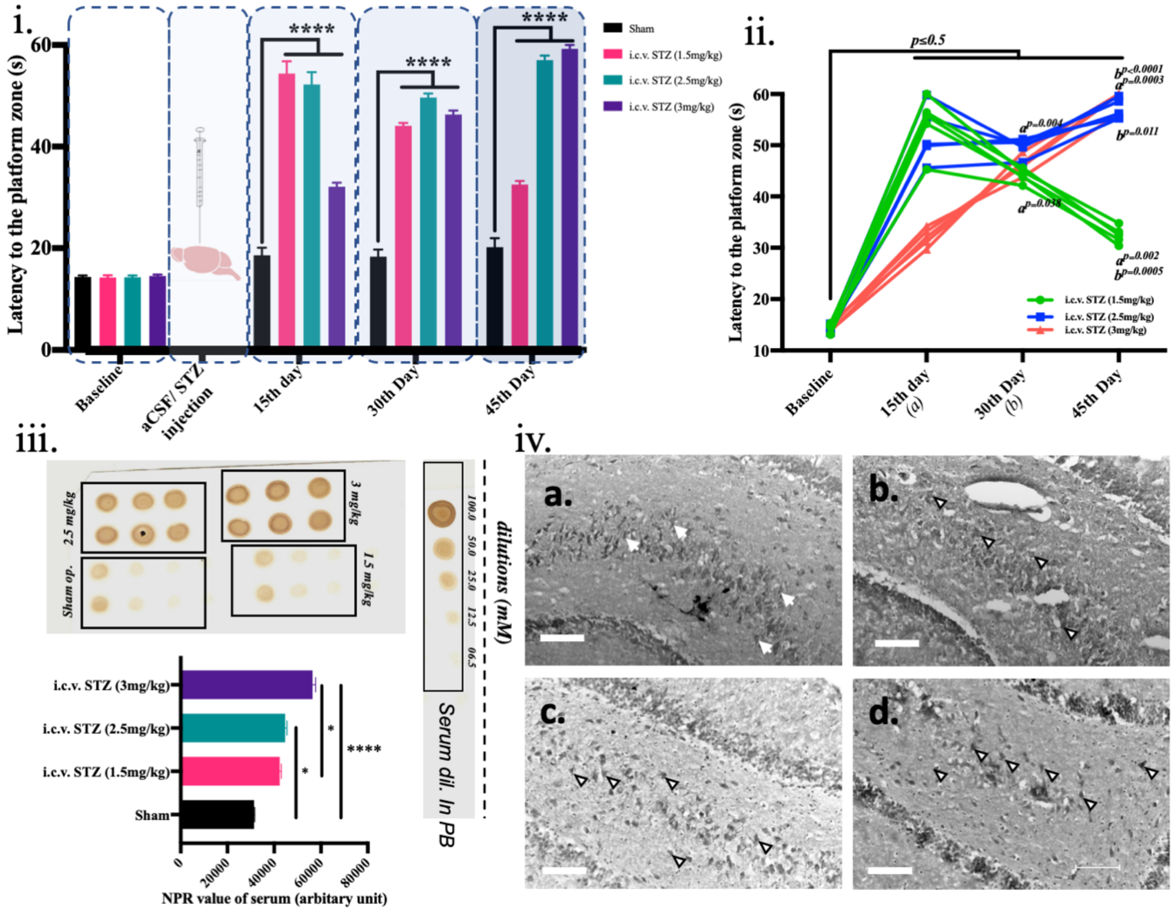


**Figure S1.b.: Selection of the streptozotocin dose for the present study.**

Latency to the platform zone at different time point (i); spaghetti line plot to compare between three doses at different time points post injection of STZ (ii); nitrocellulose redox permanganometry of the serum samples of different group of animals at 45$^{th}$ day post injection of STZ (iii); representative photomicrograph of cresyl violet stained CA3 subfield of hippocampus of sham (iv.a) 1.5mg/kg STZ (iv.b), 2.5mg/kg STZ (iv.c), 3mg/kg STZ (iv.d) injected rats; data represented as mean±SEM; n=5/group in the MWM test; n=6 biological replicate from 2 animal serum in NPR and n=3/group in cresyl violet staining group; *indicate significant where P<0.05 and ****indicate p<0.0001.

# S1.c.

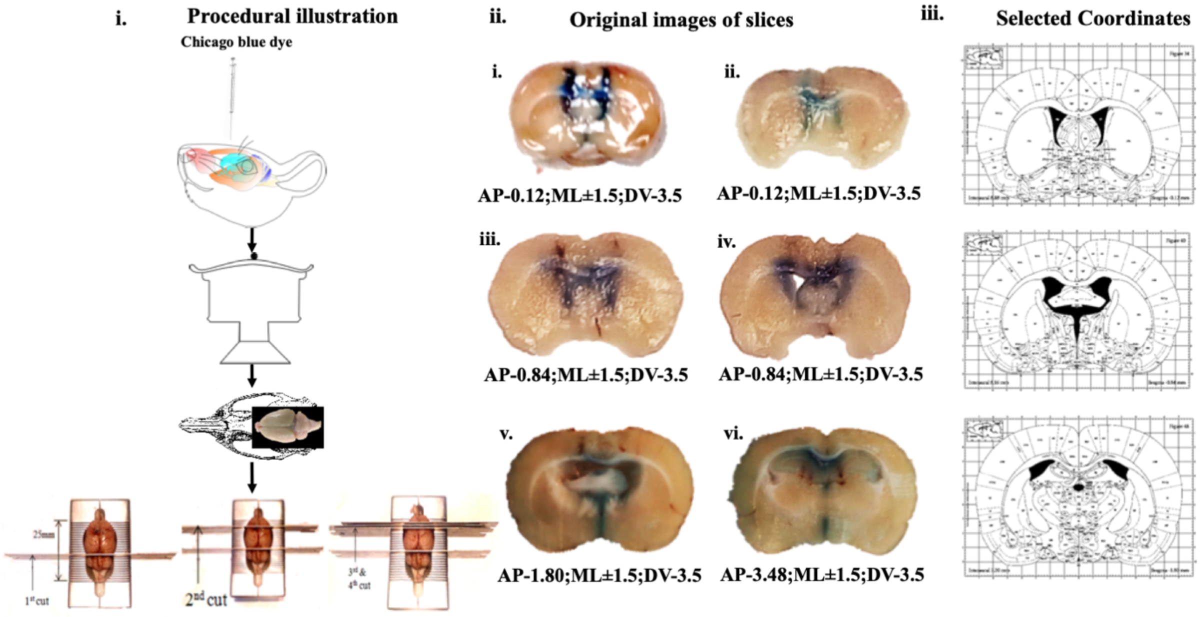


**Figure S1.c.: Selection of the stereotaxic coordinate for ICV injection of STZ.**
Procedure for injection of Chicago blue dye to the isolation of brain and 1mm section in brain matrix (i); images of the 1mm slices of dye injected brain with coordinates (ii); coordinates selected for injection of dye (iii).

**S1.d.**

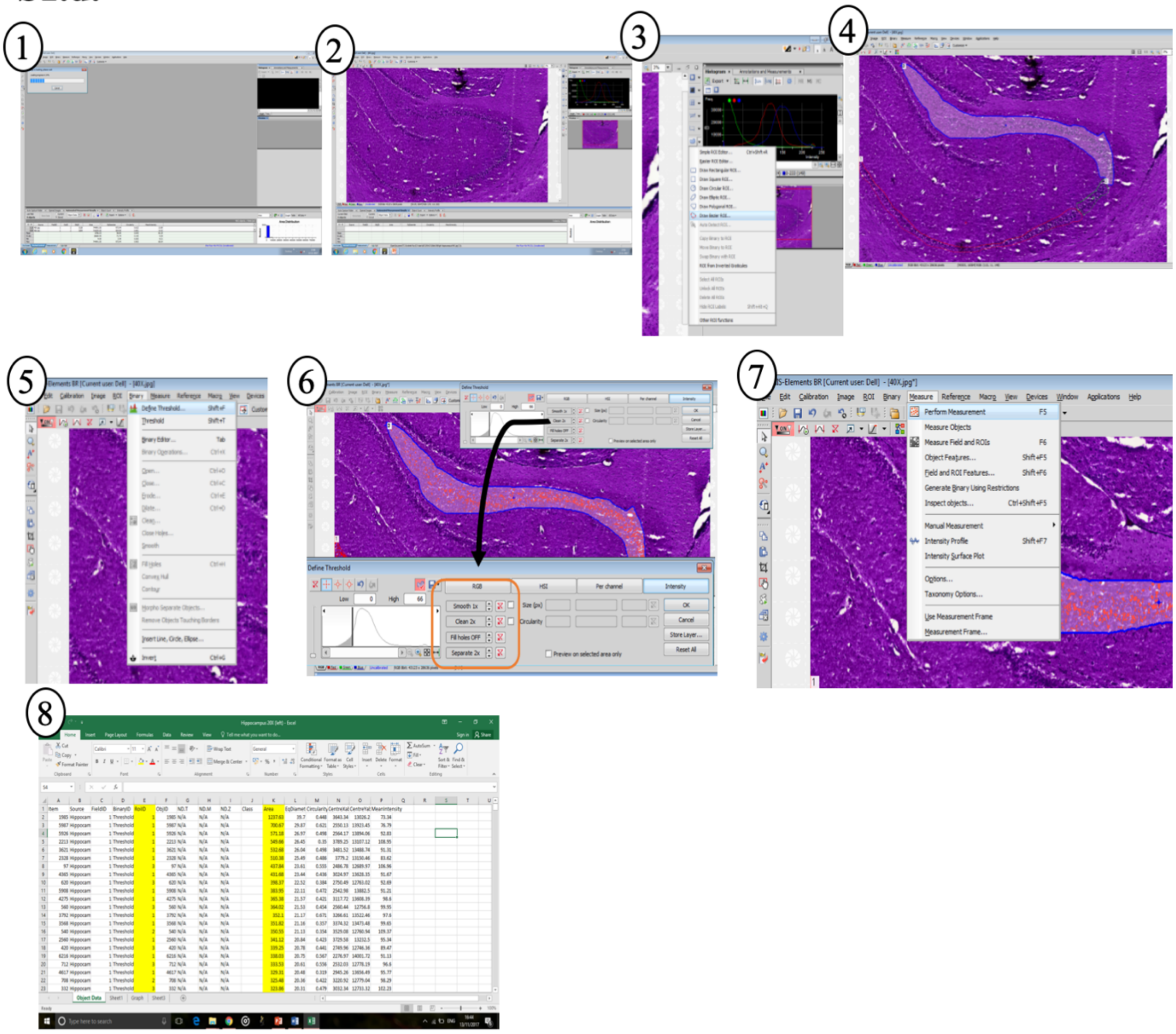

**Figure S1.d.: Workflow of counting viable cells in NIS element basic research software.** Stepwise workflow of the software with loading the image (1-2); selecting beizier drawing tool (3); drawing of region of interests (4); selection of the cells on the basis of thresholding process (5); smoothing the data using different filters (6); automatic counting using perform measurement tool (7); excel sheet including the results (8).

**Reference:**


Ennaceur, A., Cavoy, A., Costa, J.C., Delacour, J., 1989. A new one-trial test for neurobiological studies of memory in rats. II: Effects of piracetam and pramiracetam. Behav. Brain Res. https://doi.org/10.1016/S0166-4328(89)80051-8

Ferry, B., Gervasoni, D., Vogt, C., 2014. Stereotaxic neurosurgery in laboratory rodent: Handbook on best practices, Stereotaxic Neurosurgery in Laboratory Rodent: Handbook on Best Practices. https://doi.org/10.1007/978-2-8178-0472-9

Homolak, J., Kodvanj, I., Babic Perhoc, A., Virag, D., Knezovic, A., Osmanovic Barilar, J., Riederer, P., Salkovic-Petrisic, M., 2022. Nitrocellulose redox permanganometry: A simple method for reductive capacity assessment. MethodsX 9. https://doi.org/10.1016/J.MEX.2021.101611
